# Experimental evolution of gene essentiality in bacteria

**DOI:** 10.1101/2024.07.16.600122

**Authors:** Liang Bao, Zan Zhu, Ahmed Ismail, Bin Zhu, Vysakh Anandan, Marvin Whiteley, Todd Kitten, Ping Xu

## Abstract

Essential gene products carry out fundamental cellular activities in interaction with other components. However, the lack of essential gene mutants and appropriate methodologies to link essential gene functions with their partners poses significant challenges. Here, we have generated deletion mutants in 32 genes previously identified as essential, with 23 mutants showing extremely slow growth in the SK36 strain of *Streptococcus sanguinis*. The 23 genes corresponding to these mutants encode components of diverse pathways, are widely conserved among bacteria, and are essential in many other bacterial species. Whole-genome sequencing of 243 independently evolved populations of these mutants has identified >1000 spontaneous suppressor mutations in experimental evolution. Many of these mutations define new gene and pathway relationships, such as F1Fo-ATPase/V1Vo-ATPase/TrkA1-H1 that were demonstrated across multiple *Streptococcus* species. Patterns of spontaneous mutations occurring in essential gene mutants differed from those found in wildtype. While gene duplications occurred rarely and appeared most often at later stages of evolution, substitutions, deletions, and insertions were prevalent in evolved populations. These essential gene deletion mutants and spontaneous mutations fixed in the mutant populations during evolution establish a foundation for understanding gene essentiality and the interaction of essential genes in networks.

## Introduction

All living organisms require certain physical properties and biochemical capacities encoded by "essential" genes to sustain basic cellular activities. Essential genes are indispensable for an organism to survive or reproduce, and their essentiality is highly dependent on the environment in which the organism lives. ^1–6^ *In vivo*, essential gene products carry out these tasks by interacting with other components. Therefore, the essentiality of genes can also depend on the genetic context provided by the additional genes present. ^2,7,8^ Relationships may be characterized by negative epistasis when the essentiality of a gene is dependent on the absence or impairment of another gene’s function, such as synthetic lethality ^7,9^ or positive epistasis interactions, where an individual with two genes mutated is fitter than a strain possessing a mutant version of only one of the two genes. ^7,10,11^ In contrast to binary classification, it is increasingly acknowledged that essential genes exhibit a quantitative spectrum, displaying a gradient of essentiality ^2,12^, and a conditional nature. ^13^ For some essential genes, under optimal controlled conditions, deletion can still result in viable, albeit severely compromised individuals. ^14,15^ The loss of essential genes produces intense selective pressure for the fixation of genomic suppressor mutations in the descendants of the original mutants during subsequent generations, including aneuploidy of chromosomes. ^14,15^ Furthermore, essential genes can lose their essentiality during long-term evolution and acquire novel functions. ^16–18^ For instance, unlike the essential characteristics of the critical cell cycle regulator *CDK1* in yeasts and animals, ^19^ plant PSTAIRE-type cyclin-dependent kinases, known as *CDKAs* (plant homologs of *CDK1*), play a role in environmental responses that is independent of the cell cycle, ^16^ likely after the acquisition of the plant-specific cyclin-dependent kinases, *CDKBs*. ^20^

Identifying genes that interact with essential genes can produce many potential benefits, both from a theoretical perspective, such as understanding the basic principles of cellular life,^21,22^ and from a practical standpoint, such as addressing various challenges in synthetic biology ^23^ and combating drug resistance of infectious diseases. ^22^ Experimental evolution or laboratory evolution imposes a natural selective pressure on a population to redirect the evolution towards fitness-improving phenotypes in a controlled laboratory setting. ^14,15,24–30^ This methodology has been applied to identify spontaneous mutations associated with fitness-improving phenotypes, such as multicellularity, ^24^ novel metabolic capacity, ^25^ restoration of flagellar mobility,^31^ and fitness of minimal cells. ^26^ Recently, experimental evolution using mutants deleted of essential genes in the yeast *Saccharomyces cerevisiae* revealed a rapid growth phenotype after passages. ^14,15^ However, aneuploidy of chromosomes was prevalent among evolved populations deleted of essential genes, which hindered the identification of causal suppressors. ^14,15^ Here, we have established a transformation system to obtain numerous essential gene knockout mutants, including *obgE*, *pfkA*, components of Type-II fatty acid synthesis (FAS II) and subunits of F1Fo-ATPase genes that have been identified as essential in most bacterial species, including all species of *Streptococcus* examined (http://www.essentialgene.org/). Therefore, most of these mutants represent the first of their kind in the entire streptococcal field. For the 23 essential genes whose deletion resulted in a slow-growing phenotype, we performed experimental evolution of their corresponding mutants. After allowing for short-term adaptation, we have identified >1000 spontaneous suppressor mutations in 243 evolved populations deleted of essential genes, most of which are substitutions, deletions, and insertions. This feature enabled us to map virtually all mutations to distinct genomic segments of individual ORFs or intergenic regions. As a proof of concept, we have demonstrated the *f1fo* genes within the F1Fo-ATPase/V1Vo-ATPase/TrkA1-H1 gene pathway across multiple Streptococcus species. For comparison, we also evaluated the twelve mutations that arose in six evolved populations in the wild-type (WT) cells. The mutations that occurred in essential gene mutants displayed several characteristics that distinguished them from those that evolved in WT cells during short-term evolution.

## Results

### Isolating essential gene mutants and analysis of short-term evolution

The genome of *Streptococcus sanguinis* SK36 comprises over 2000 ORFs, ^32^ of which 218 have been experimentally identified as essential for the organism’s survival when cells are grown on a rich medium—brain heart infusion (BHI) broth or plates—under microaerobic conditions (Fig. 1a). ^1^ To isolate viable essential-gene deletion mutants, we have modified our transformation procedure to minimize stress (see Materials and Methods). Although we still employed homologous recombination (Fig. 1b and Supplementary Fig. 1a), we replaced our conventional 1-hour aerobic transformation with an anaerobic incubation and a duration of up to 24 hours. ^1^ Additionally, we optimized the selection conditions by covering the transformants with a thin layer of agar medium and extending the selection period for the transformants to four to six days, all conducted under anaerobic conditions. (See Materials and Methods.)

**Figure 1.**
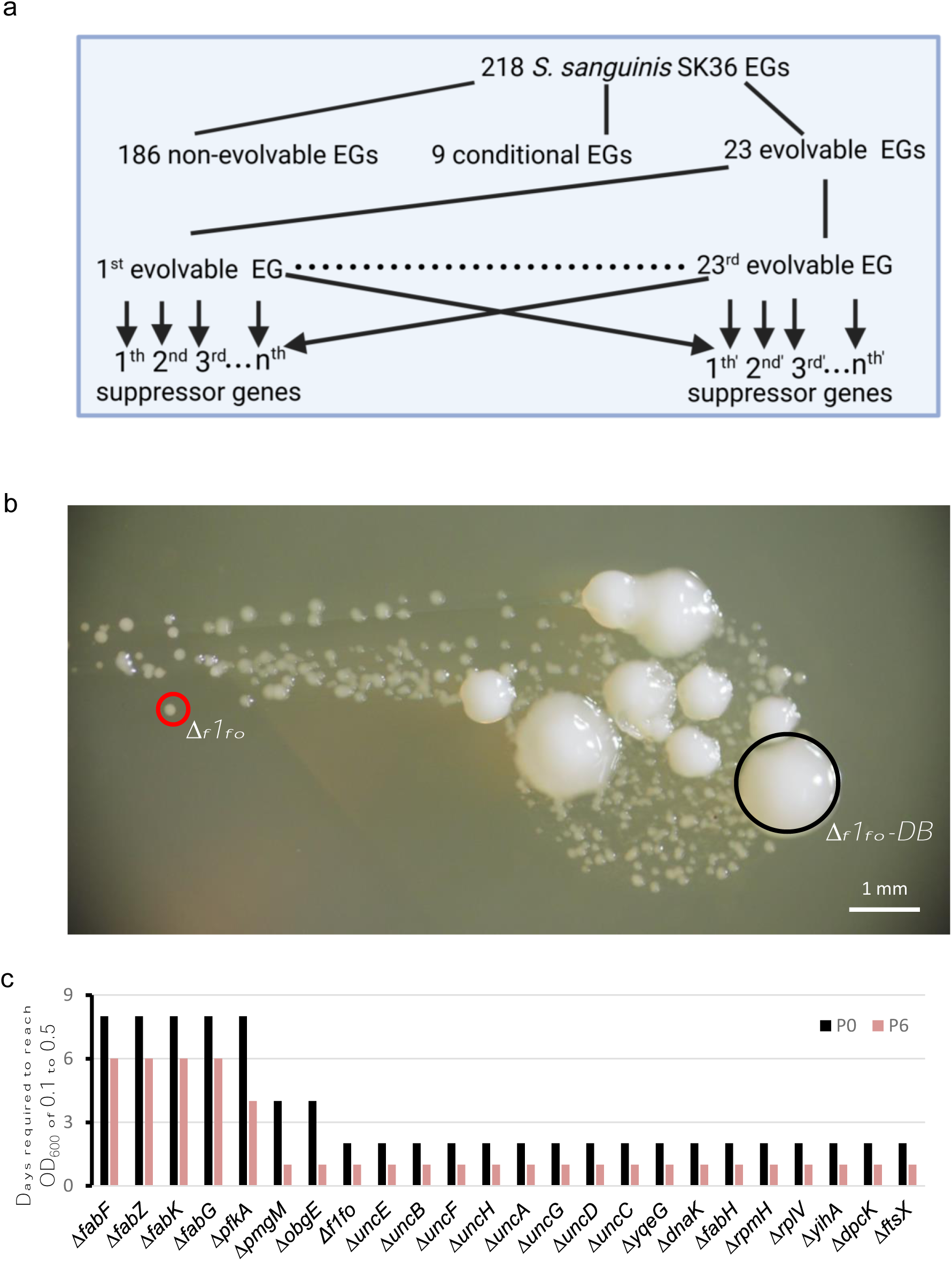
Evolution of mutants deleted for essential genes in passage experiments. (a) The 218 essential genes (first row) were categorized into three groups: 186 non-evolvable essential genes (EGs), 9 conditional EGs, and 23 evolvable EGs (second row). Knockout mutants of the evolvable EGs (third row) were subjected to experimental evolution to identify suppressor genes (fourth row) in the mutant populations. Arrow headed lines indicate the suppressor genes that evolved in populations deleted of evolvable EGs. (b) Two types of colonies, small and large, as demonstrated for *f1fo* deletion, appear on selection plates after 5 days of growth. Only the small colonies represent true deletions, denoted as *Δf1fo* (red circle). The large colonies are "double-band mutants," containing both the replacement of the original *f1fo* with a *kan* gene and a wild-type *f1fo* copy, or they are mutants that did not contain a *kan* gene, but contained point mutations in other genes, denoted as *Δf1fo-DB* (black circle, ’DB’ for double-band). (c) Incubation time (days) required for P0 and P6 mutants to reach an OD_600_ of 0.1 to 0.5.

Using this procedure, we observed an interesting phenomenon. Specifically, for 23 of the essential genes, the transformants produced two types of colonies: large and small (Fig. 1a, 1b, and Supplementary Fig. 2a). As expected, genotyping of the small colonies revealed a complete replacement of the corresponding ORF with a kanamycin resistance (*kan*) gene (Supplementary Fig. 1b). In contrast, genotyping of the large colonies showed that some mutants displayed a "double band" genotype due to duplication of the target gene prior to the replacement of one copy by the *kan* gene (Supplementary Fig. 1c, Supplemental Table 1). This finding suggests that *S. sanguinis* SK36 regularly undergoes gene duplications, particularly between two copies of a multi-copy gene or locus, such as the rRNA operons. Alternatively, some large colonies lacked a *kan* gene, but contained point mutations in different genes, with the majority of isolates acquiring mutations in translational machinery genes that may confer resistance to kanamycin, ^33^ such as *fusA, rpsL* or *rplF* (Supplemental Table 1). Among fifty-four large colonies genotyped, fifteen showed the double-band genotype. The remaining thirty-nine did not contain the *kan* gene, but thirty-eight, or 97.4% contained mutations in *fusA, rpsL* or *rplF* (Supplemental Table 1). In contrast, for nine essential genes, the deletion mutants exhibited robust growth (Supplementary Fig. 2b). These nine genes included *nrdE, nrdF, nrdH,* and *nrdI* which were previously reported as non-essential under anaerobic conditions but essential in the presence of O_2_, ^34^ along with *rexB* for ATP-dependent nuclease subunit B, *rpsA* for 30S ribosomal protein S1, *pdf* for peptide deformylase, and *g6pD* for glucose-6-phosphate dehydrogenase (Fig. 1a, Supplementary Fig. 2b).

We, therefore, directed our attention to the twenty-three essential genes that, when deleted, led to severe growth defects in the resultant mutants, denoted as evolvable essential genes (Fig. 1a). ^14,15^ These twenty-three genes encoding components of diverse pathways, include the eight subunit genes encoding the F1Fo-ATPase (*uncE*, *uncB*, *uncF*, *uncH*, *uncA*, *uncG*, *uncD*, and *uncC)*; *ftsX,* encoding a permease-like cell division protein; five *fab* genes (*fabH*, *fabK*, *fabG*, *fabF*, and *fabZ*) that are involved in Type-II fatty acid synthesis (FAS II); two ribosomal protein genes (*rplV* and *rpmH*); two GTPase genes (*obgE* and *yihA*); *yqeG,* encoding a YqeG-family phosphatase; *pfkA*, encoding 6-phosphofructokinase, involved in glycolysis; *pngM*, encoding phosphoglucomutase; *dpcK* (also called *coaE*), encoding diphospho-CoA kinase, involved in CoA synthesis; and *dnaK*, encoding a heat shock protein chaperone. These genes are widely distributed and are essential in various genera and species (http://www.essentialgene.org/). ^35^ In total, we have created 23 essential gene mutants, each with an individual gene deletion, along with one mutant deleted for the entire *f1fo* region encoding the eight F1Fo-ATPase subunits, which we have named *Δf1fo* (Supplementary Fig. 1 and S2a.1-a.2). Although the deletion mutants of the 23 genes were initially generated under anaerobic conditions in this study, we have confirmed the viability of mutants under both anaerobic (Supplementary Fig. 3a) and microaerobic (Supplementary Fig. 3b) conditions, as utilized previously for identification of essential genes in *S. sanguinis* SK36. ^1^ To further verify that the viability of these 24 gene deletion mutants was not due to the absence of oxygen, we randomly selected three essential gene deletions—*pfkA*, *dpcK*, and the entire *f1fo* region—and confirmed that we were able to obtain viable, though extremely slow-growing, target gene deletion mutants under microaerobic conditions as well (data not shown).

In order to study the evolution of gene essentiality, we conducted passage experiments using multiple independently evolved populations of the 24 essential gene mutants. To form a population, we selected three to five colonies from the selection medium, combined them into a single inoculum, and subjected them to passage in BHI by 1:20 serial transfer. To ensure a comparable number of generations, we closely monitored the growth of mutants during each passage. We proceeded to the following passage only when the cell cultures had reached an OD_600_ within the range of 0.1-0.5. As a control, WT cells, which reached an OD_600_ of approximately 1.0 within 24 hours of growth, or 2-10 times more cells compared to the mutants in each passage, were passaged daily for a total of nine times (Fig. 1c). This method was applied to maintain consistency throughout the experimental process. While all mutants exhibited significant growth defects compared to WT, we observed variations in their growth phenotypes. Notably, mutants lacking *pfkA, fabK, fabG, fabF*, and *fabZ* displayed the most severe growth defects. In contrast, the growth of others, including another *fab* mutant, *fabH,* was less severe, as evidenced by the OD_600_ measurements of the evolved mutants (Fig. 1c and Supplementary Fig. 3c, Supplemental Table 2). In our passage experiments, we observed that the mutants in passage 6 (P6, "P" for passage) exhibited enhanced fitness compared to those in early passages. For example, the time required for cultures to reach an OD_600_ of 0.1-0.5 in P0 compared to P6 (Fig. 1c, Supplemental Table 3) indicated faster growth at P6, reflecting improved fitness.

### Identification of suppressor loci in evolved populations deleted of essential genes and in WT

To identify suppressor mutations in the evolved populations deleted of essential genes, we performed whole-genome sequencing at P6 (Fig. 1c). Specifically, we sequenced between two and twenty-five independently evolved populations for each of the twenty-four essential gene mutants, resulting in a total of 243 populations (Fig. 2a and Supplemental Table 4). As controls, we also sequenced six populations of WT, which were generated by aliquoting into six separate tubes and passaging nine times, as well as the original WT (Fig. 2a and Supplemental Table 4).

**Figure 2.**
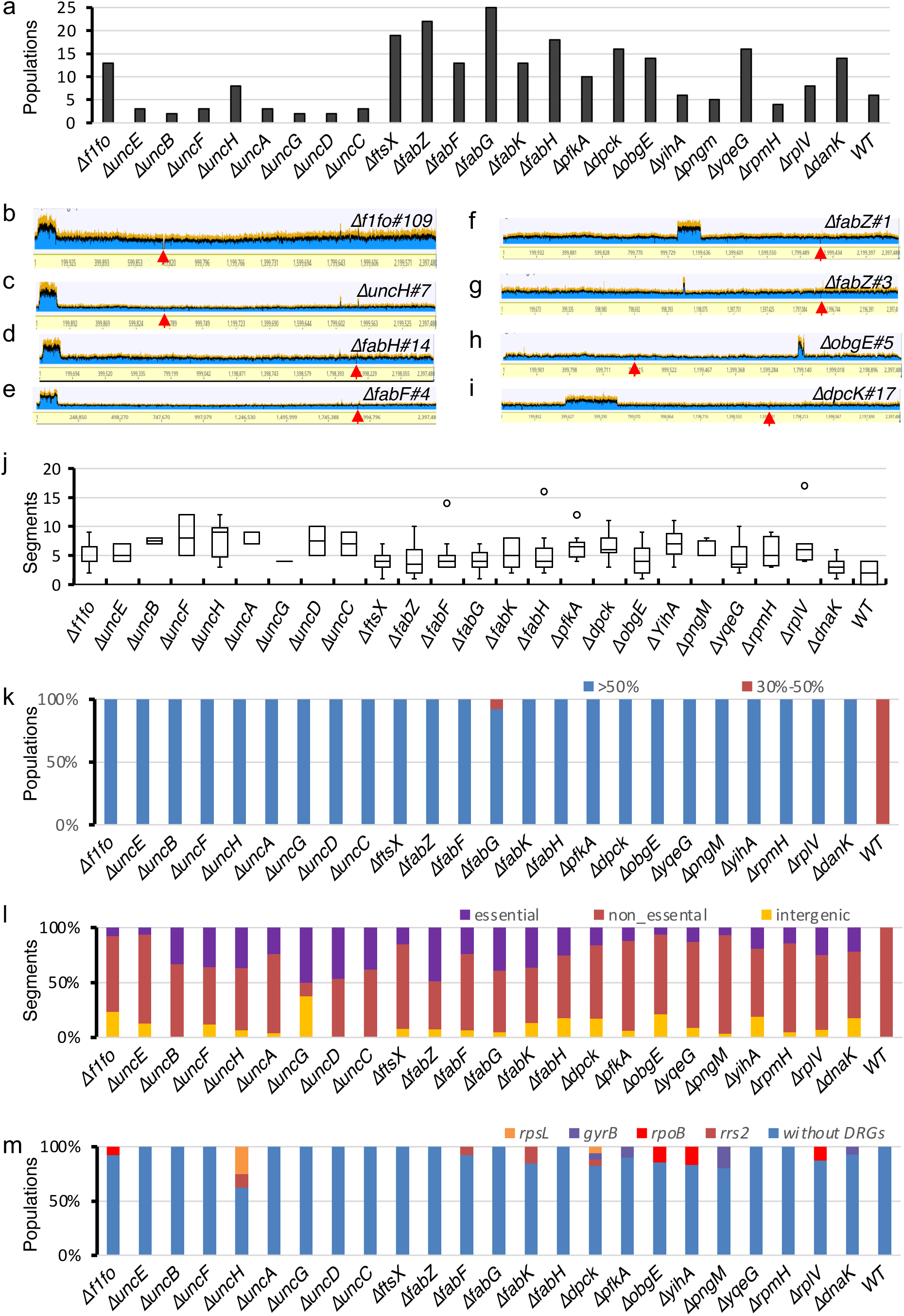
Identification of suppressors in mutants deleted for essential genes. a) Number of independently evolved populations of different mutants. (b-i) Evolved populations containing large-scale duplications. Red arrow, site of original essential gene deletion; Y axis indicates number of sequence reads mapped to the reference sequence at the coordinates shown on X axis. (j) Boxplot of mutated segments (≥30% abundance) in the independently evolved populations of mutants deleted for essential genes or WT. Open circles indicate outliers. (k) Stacked proportions of populations containing mutated segments comprising >50% (blue) or 30%-50% (red) of sequence reads. (l) Stacked proportions of mutated segments belonging to intergenic regions, non-essential ORFs, or essential ORFs. (m) Stacked proportions of evolved populations with/without mutations in the multi-drug resistance genes (DRGs: *rpsL*, *gyrB*, *rpoB* and *rrs2*) shown.

We first considered mutations related to duplications of genes other than the essential gene targeted for mutagenesis. Of the 243 evolved mutant populations, we found that twelve (4.9%) displayed gene duplications of various sizes, ranging from 10 kb to 310 kb. By comparison, none of the six evolved WT populations in P9 nor the original parental WT contained a gene duplication (Supplementary Fig.4a). For the twelve evolved mutant populations with gene duplications, two were from *Δf1fo* (*#107* and *#109*) (Fig. 2a and Supplementary Fig.4b), one from *ΔuncE* (*#2*) (Supplementary Fig.4c), one from *ΔuncH* (*#7*) (Fig. 2c), three from *ΔfabH* (#4, #6 and #14) (Fig. 2d and Supplementary Fig.4d and S4e), one from *ΔfabF (#4)* (Fig. 2e), two from *ΔfabZ* (*#1 and #3*) (Fig. 2f and 2c), one from *ΔobgE* (*#5*) (Fig. 2h) and one from *ΔdpcK* (*#17*) (Fig. 2i). The duplicated regions found in populations of *Δf1fo, ΔuncE, ΔuncH, ΔfabH* and *ΔfabF* mutants encompassed the same ∼103 kb region (coordinate: 22,738-125,834) that is flanked by two directly repeated rRNA operons (Fig. 2b-e). The above-duplicated region of ∼103 kb contains 113 ORFs, including nine ORFs encoding the entire V1Vo-ATPase operon that is related to the proton pump function of F1Fo-ATPase, ^36^ one ORF encoding acyl carrier protein and one ORF encoding acyltransferase that is related to lipid biosynthesis. ^37^ The duplicated region found in *ΔfabZ#1* spans approximately 143 kb (coordinate: 1,053,488-1,197,391). The 10.1-kb duplication in *ΔfabZ#3* (coordinate: 1,093,824-1,103,958) is within the duplicated 143 kb in *ΔfabZ#1.* This 10.1-kb region contains ten ORFs, including one ORF annotated as acyl-acyl carrier protein thioesterase, which catalyzes the terminal reaction of fatty acid biosynthesis (Jing, et al., 2011). In the case of *ΔobgE#5*, a 37-kb duplication is present (coordinate: 1,775,394-1,812,805), encompassing thirty ORFs, including one ORF encoding der GTPase. In *ΔdpcK#17,* a ∼310-kb duplication occurred (coordinate: 385,678-697,581), encompassing 305 ORFs. The flanking sequences of the duplicated regions in *ΔfabZ#1, ΔfabZ#3, ΔobgE#5* and *ΔdpcK#17* possessed no notable repeated sequences.

Next, we assessed substitutions, deletions, and insertions of less than 10 kb in the evolved populations. The genome of *S. sanguinis* can be divided into 4,283 segments—2,340 ORFs and 1,943 intergenic regions (Supplemental Table 5). ^38^ We defined a segment as mutated if any mutation within that segment was present in at least 30% of the sequence reads (see Materials and Methods section). We selected 30% as the threshold based on our finding that confirmed suppressor mutations, such as *trkA1-H1*, ranged from 34.8% to 100% in the evolved *Δf1fo* populations in P6 (Fig. 4d and Supplemental Table 4). There were six mutations (two substitution mutations and four deletion mutations from the six evolved populations), where two to four ORFs were affected by each mutation. For example, the C to T (at coordinate 1,313,563) mutation in *ΔfabF#10* caused amino substitutions of two overlapping ORFs encoding two hypothetical proteins (Supplemental Table 4). The remaining mutations fell within single segments of ORFs or intergenic regions. In total, we detected 1,272 mutated segments (Supplemental Table 4), with 1,260 from evolved populations lacking essential genes and twelve from the evolved WT populations (Supplemental Table 4). Our analysis revealed that each of the 243 (100%) evolved mutant populations contained at least one mutated segment, with the majority containing from 2 to 10 (Fig. 2j and Supplemental Table 4-6). In contrast, among the six evolved WT populations, four had at least one segment containing a mutation in at least 30% of the sequence reads, while the remaining two, as well as the parental WT population, had none (Fig. 2j and Supplemental Table 4). Furthermore, when we set the threshold to 50%, 241 evolved mutant populations, or 99.2%, had mutated segments, whereas none of the six evolved WT populations contained mutated segments (Fig. 2k and Supplemental Table 7) (χ2=154.69, df=1, n=249 populations, p=2.2e-16).

We then analyzed the presence of mutated segments belonging to essential ORFs (in each case, excluding the ORF that was intentionally deleted in each mutant, as well as the nine conditionally essential genes). We found that suppressor mutations belonging to essential segments were present in at least one evolved population from each essential gene mutant (Fig. 2l and Supplemental Table 8-9). In contrast, none of the twelve mutated segments from the evolved WT populations were found to be within an essential ORF (Fig. 2l and Supplemental Table 8-9) (χ2=5.74, df=1, n=25 groups, p=0.017). To illustrate this further, we compared the frequency of mutated segments in P1 and P6 for two specific cases: mutations of *ftsZ* (an essential segment encoding a structural homolog of tubulin in prokaryotes) in the evolved *Δf1fo#107* and *uncH* (an essential segment, encoding delta subunit of F1Fo-ATPase) in the evolved *ΔuncC#1*. The *uncH* gene is located 3.9 kb upstream of *uncC*, both of which are within the *f1fo* operon. The evolved *ΔuncC#1* population of P1 did not have any *uncH* mutations using 30% as the cutoff, although an *uncH* mutation was present in 12% of the sequence reads (Supplementary Fig. 5a-c). In contrast, the *ΔuncC#1* population of P6 contained 86% *uncH* mutation (Supplementary Fig. 5d-f) The resultant mutation replaced a 207-bp segment that was 51 bp downstream of the *uncH* start codon with an 82-bp fragment, causing early truncation (Supplementary Fig. 5g). For the second case, the *ftsZ* gene is located approximately 126 kb upstream of the *f1fo* operon. In the evolved *Δf1fo#107* population in P1, we did not detect any mutations in the *ftsZ* segment using 30% abundance as cutoff, although 27% of the sequence reads contained a *ftsZ* mutation (Supplementary Fig. 5g-h). In comparison, the evolved population of P6 contained 100% mutated *ftsZ*—a 248 bp insertion located 1015 bp downstream of the start codon (Supplementary Fig. 5i-j), leading to the truncation of the FtsZ protein (Supplementary Fig. 5k). These findings demonstrate an increase of mutated essential segments in the essential gene deletion mutant due to selection during passage.

Multi-drug resistance across ESKAPE pathogens (i.e., *Enterococcus faecium*, *Staphylococcus aureus*, *Klebsiella pneumoniae*, *Acinetobacter baumanii*, *Pseudomonas aeruginosa*, and *Enterobacter spp.*) has been found to be commonly associated with mutations in six genes— namely, *rrs, gyrA*, *gyrB*, *rpoB, rpsL* and *rsmG* (also known as *gidB*). ^39^ We therefore asked whether mutations in the typical set of multi-drug resistance-associated genes ^39^ could appear in the evolved populations deleted of essential genes. Apart from the 16S rRNA gene, which is present in four copies (*rrs1* to *rrs4*), *S. sanguinis* has one copy of each of these genes. Among the 243 evolved populations deleted of essential genes, mutations of multi-drug resistance genes were present in *rrs2*, *rpoB*, *gyrB,* and *rspL* in a variety of populations derived from different essential gene mutants, while no mutations were detected in *rrs1*, *rrs3*, *rrs4*, *gyrA* or *rsmG*. Moreover, no mutation was detected within this set of multi-drug resistance-associated genes in the evolved WT populations (Fig. 2m and Supplemental Table 10). These results showed that missense mutations in four of the same genes that promote multi-drug resistance also improve the fitness of mutants deleted of these particular essential genes.

In summary, compared to the evolved WT populations, mutants deleted for essential genes exclusively evolve gene duplications of greater than 10 kb, mutations in essential genes, and mutations of genes associated with multi-drug resistance. In addition, once mutations are observed, they are more quickly fixed in the population (Fig. 2k).

### Construction of a network using WT and essential gene deletion mutants and their suppressors

To explore the relationships of suppressor mutations from the evolved populations, we established a network of 25 groups, comprising the 24 mutants deleted for essential genes, along with SK36, and their respective evolved suppressors across 249 populations (243 populations from mutants and six populations from SK36). In these 249 populations, 1,272 mutations occurred in 592 distinct segments, with 503 segments belonging to ORFs and 89 belonging to intergenic regions (Fig. 3a, Supplemental Table 4, and Supplemental Table 11-13). This network also encompasses a total of 914 edges connecting these 617 nodes (Fig. 3a and Supplemental Table 11-13).

**Figure 3.**
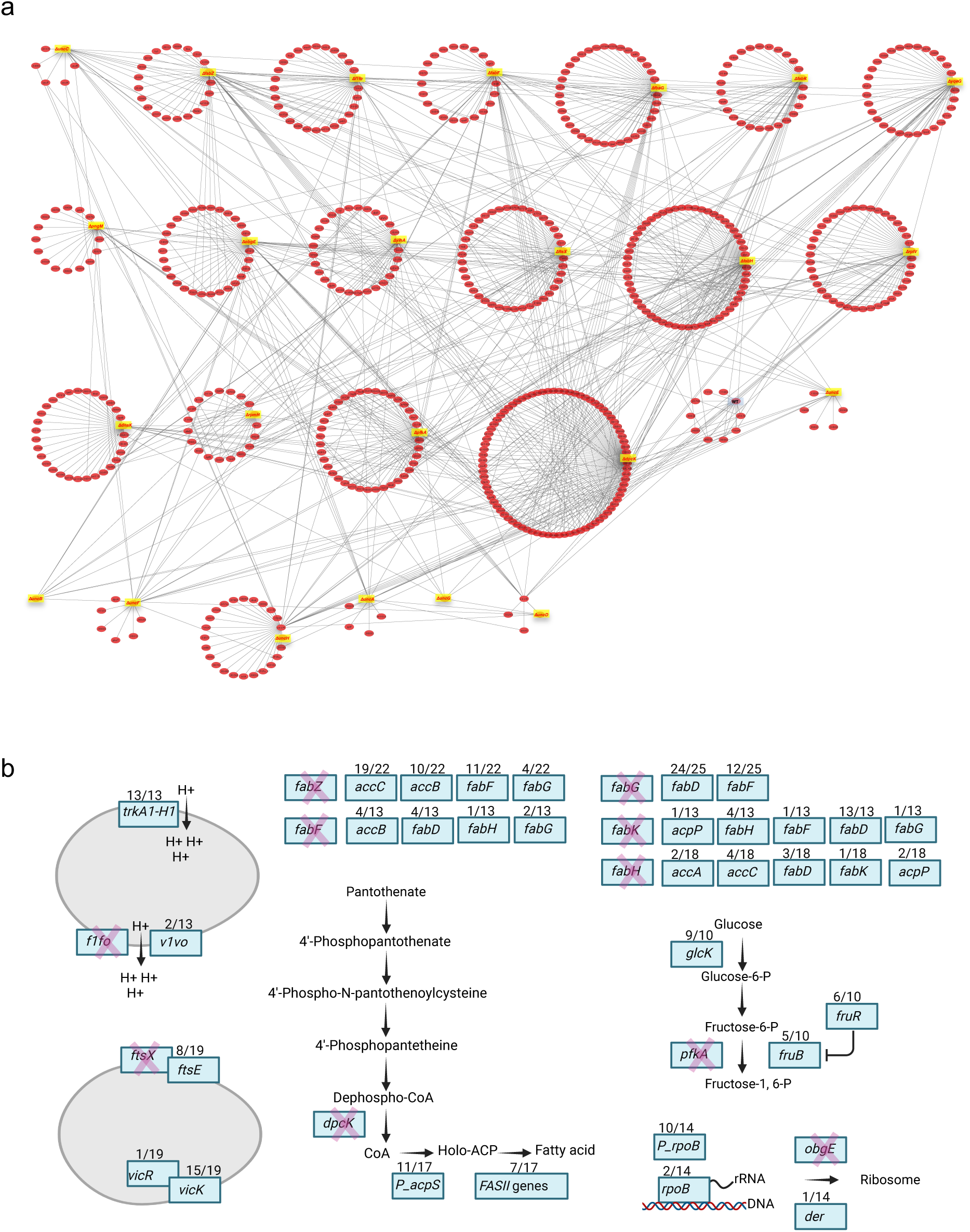
Networks derived from the mutations in the evolved populations. (a) A network was built from the lists of suppressor mutants derived from each essential gene mutant and WT. Yellow-highlighted labels indicate the names of the essential genes mutated and the blue-highlighted label indicates the passaged WT. In total, there are twenty-five groups. Gene names without any marks, mutated segments belonging to essential genes; gene names with an asterisk (*) on the left, conditional essential genes; gene name with two asterisks on the left, non-essential genes; P_ followed by gene names and an asterisk, promoters; In_ followed by gene names and an asterisk and int_ followed by number, intergenic regions; _ followed by number, gene locus. (b) Most commonly mutated segments in the evolved populations deleted of essential genes. Red cross on the rectangle boxes indicates evolved populations deleted of essential genes. Labels within the rectangle boxes indicate the segments with mutations. Numbers on top of the rectangle boxes indicate number of populations with mutations in the segments out of the total populations. Arrows indicate the flow of biochemical reactions.

Some mutations discovered in the evolved WT also appeared in evolved mutant populations. The twelve mutations occurred in ten distinctive segments, with 5 segments specific to WT and 5 shared with populations derived from essential gene mutants. Notably, the ORF J1C87_09360, annotated as PTS mannose/fructose/sorbose transporter family subunit IID, exhibited mutations in two of the six evolved WT populations. This result is consistent with a previous study in SK36, where fitness improvement was associated with mutations in three ORFs of the same operon (J1C87_09355, J1C87_09360, and J1C87_09365). ^40^ In the current study, mutations in the three OFRs were also observed in mutant populations of *ΔuncH, ΔfabG, ΔfabK, ΔfabF, ΔfabZ, and ΔpfkA* (Supplementary Fig. 6a-b and Supplemental Table 14). Interestingly, most of the evolved populations that contain suppressor mutations in the three PTS mannose/fructose/sorbose transporter subunits are also those that showed the most severe growth defects, such as *ΔfabG, ΔfabK, ΔfabF, ΔfabZ, and ΔpfkA* (Fig. 1d and Supplementary Fig. 3c). These results demonstrate that mutations in some segments that apparently improved fitness in WT could also contribute to the fitness improvement of essential gene mutants.

Mutations in some segments that were present in evolved populations deleted of one essential gene also appeared in evolved populations of other mutants deleted of another essential gene. For example, most populations of the mutant deleted for the entire *f1fo* region (*Δf1fo*) and the eight F1Fo subunits deletion mutants (*ΔuncE, ΔuncB, ΔuncF, ΔuncH, ΔuncA, ΔuncG, ΔuncD* and *ΔuncC)* contain suppressor mutations in J1C87_08610 (*trkA1*) and J1C87_08615 (*trkH1*) (Fig. 3a). Mutations in many F1Fo subunits are shared among the evolved populations of various other subunit deletion mutants (Fig. 3a and Supplemental Table 4). Furthermore, many of the suppressor mutations in type II fatty acid synthesis (FAS II) gene mutants, such as *ΔfabZ, ΔfabF, ΔfabG, ΔfabK* and *ΔfabH* and those in the CoA biosynthesis mutant *ΔdpcK*, occur in the same ORFs related to fatty acid biosynthesis, including essential genes like acetyl-CoA carboxylase carboxyl transferase subunit alpha (J1C87_09410: *accA*) and acetyl-CoA carboxylase biotin carboxylase subunit (J1C87_09420: *accC*) (Fig. 3a). Similarly, the suppressors identified in the populations of two ribosomal protein mutants *(ΔrpmH* and *ΔrplV*) and two GTPase mutants (*ΔyihA* and *ΔobgE*) include mutations in 50S ribosomal protein gene *rplB* (J1C87_00620) and RNA biosynthesis genes, such as *rpoB* (J1C87_01040) (Fig. 3a and Supplemental Table 4). These findings indicate that mutated segments shared across multiple evolved mutants can confer fitness benefits to mutants deleted of different essential genes.

We also observed mutual suppressors, where an evolved population deleted of the first essential ORF contained a mutated segment within a second essential ORF, while an evolved population deleted of the second essential ORF contained a mutated segment within the first essential ORF. For example, in the *obgE* deletion mutant *(ΔobgE*), a mutation in *dnaK* occurred (Fig. 3a and Supplemental Table 4); conversely, in *ΔdnaK*, a mutated segment of *obgE* was detected (Fig. 3a and Supplemental Table 4). Similarly, in the F1Fo subunit deletion mutant *ΔuncF*, a mutation of *dpcK* was also present (Fig. 3a and Supplemental Table 4). In *ΔdpcK*, mutated segments were detected within *uncA* and *uncD* (Fig. 3a and Supplemental Table 4). Other mutual suppressor pairs include those between subunits of F1Fo (Fig. 3a and Supplemental Table 4) and between components of the FASII system (Fig. 3a and Supplemental Table 4). These results suggest the presence of positive epistasis interactions between different essential gene(s), where strains containing mutations in both essential genes are fitter than those containing mutations in either of the essential genes.

### Possible explanations for parallel suppressor evolution in mutants deleted for particular essential genes

Given the potential role of parallel evolution for fitness improvement,^41–43^ we next sought to identify the most commonly mutated segments for each essential gene mutant and those clustered in a particular operon. Utilizing these highly occurring suppressors (Supplemental Table 4 and Supplemental Table 16), along with candidate genes within duplicated regions (Fig. 2b-h), we aimed to elucidate the mechanisms by which they improved the growth of mutants deleted for essential genes.

#### Δf1fo suppressors

F1Fo is proposed to function as an ATP-dependent proton pump in streptococci, including *S. mutans* and *S. sanguinis*. ^44^ Mutations in *trkA1* or *trkH1* are present in all thirteen evolved populations of *Δf1fo*, with twelve populations containing *trkH1* mutations and one containing a *trkA1* mutation (Fig. 3b and Supplementary Fig. 7, Supplemental Table 4 and Supplemental Table 16). Similarly, mutations in *trkA1*, *trkH1*, or a one base pair (bp) insertion mutation located 38 bp upstream of the *trkA1-H1* operon (P_trkA1-H1, "P" for promoter) are found in twenty-five out of twenty-six (96.2%) of the evolved populations deleted of single F1Fo subunits (Fig. 3b and Supplementary Fig. 7, Supplemental Table 4 and Supplemental Table 16). Among the twenty-three mutations in the ORFs of *trkA1-H1*, sixteen populations have *trkH1* mutations, and eight have *trkA1* mutations. Given the comparable length of *trkA1* (1350 bp) and *trkH1* (1440 bp), it appears that mutations in the *trkH1* gene are more likely than mutations in *trkA1* to improve the fitness of mutants deleted for F_1_F_o_ subunits.

We also observed that gene duplications occurred in the same region across populations deleted of all F1Fo subunits or single subunits, i.e., *Δf1fo#107* and *#109* (Fig. 2b and Supplementary Fig. 4b), *ΔuncE#2* (Supplementary Fig.4c) and *ΔuncH#7* (Fig. 2c). This region of approximately 103 kb (coordinates: 22,738-125,834) contains the *v1vo* operon encoding the V1Vo-ATPase, which has been proposed to function as a proton pump ^36^ and may compensate for the loss of *f1fo*. It is interesting to note that *ΔuncH#7* is the only evolved population listed above that lacks mutations in the flanking upstream regions or the ORFs of *trkA1* or *trkH1* (Fig. 2c). More interestingly, among the evolved populations deleted of F_1_F_o_ subunits (thirteen *Δf1fo* and twenty-six mutants deleted for single *unc* subunit genes), *ΔuncE#2* and *ΔuncH#7* are the only two populations that contain amino acid substitution mutations of V-type ATP synthase subunits (Supplemental Table 4 and Supplemental Table 16).

Suppressor mutations appeared in other F1Fo subunits or intergenic regions of *f1fo* in many evolved populations deleted of a single subunit (Fig. 3b and Supplementary Fig.7). This result suggests that for the F_1_F_o_ complex, when one subunit is absent, other subunits cannot function properly and their presence is detrimental to the mutant.

Given that V1Vo may also function as an ATP-dependent proton pump ^36^ and the Trk system works as a symporter (K+/Na+ or K+/H+), ^45^ we hypothesize that *v1vo* can partially compensate for the loss of *f1fo*, that TrkA1-H1 is the major contributor to acidification of the cytosol when pumping in K+, and that the function of TrkA1-H1 must therefore be inhibited when F1Fo is not present (Supplementary Fig. 11).

#### ΔftsX suppressors

FtsEX is crucial in regulating divisome assembly and activation at the plasma membrane and cell wall. ^46^ In the evolved *ΔftsX* populations, mutations in the *ftsE* gene of the *ftsEX* operon and the *vicRK (vicK/walK* and *vicR/walR)* two-component system (TCS) frequently occurred (Fig. 3b and Supplementary Fig. 7). Mutations in *ftsE* were present in eight out of nineteen evolved *ΔftsX* populations (42.1%); in one of these populations, *ΔftsX#13*, a deletion mutation at 51 bp of the 693 bp *ftsE* ORF leads to a frameshift in the N-terminal region (Fig. 3b and Supplementary Fig. 7). This observation suggests that in the absence of *ftsX*, *ftsE* does not function properly and is deleterious to the cell. Although *vicR* and *vicK* compose a TCS, *vicK* is not essential, while *vicR* is essential. ^1,47^ Fifteen populations contained *vicK* mutations (with one frameshift and one truncation), and one population contained a *vicR* mutation (an amino acid substitution, M54V), collectively representing 89.5% of the evolved populations (Fig. 3b and Supplementary Fig. 7, Supplemental Table 4 and Supplemental Table 16). It was reported that FtsEX interacts with the cell wall component peptidoglycan hydrolase, PcsB, which is positively regulated by the *vicRK/walRK* two-component regulatory system. ^48^ Therefore, we hypothesize that when FtsEX is absent and PcsB cannot function properly, there is a need to inhibit the activity of PcsB’s upstream regulatory factor, VicRK.

#### fab operon suppressors

The *fabH, fabF, fabG, fabZ, and fabK* genes belong to a FASII operon containing thirteen ORFs in total, with a MarR family transcriptional regulator gene (*fabT*) upstream and seven other genes. In the evolved populations of *ΔfabZ, ΔfabF, ΔfabG, ΔfabK,* and *ΔfabH*, we observed a significant enrichment of mutations in genes encoding components of the FASII system (Fig. 3b and Supplementary Fig. 7, Supplemental Table 4, and Supplemental Table 16). This result suggests that when one component of the FASII system is deleted, this can trigger genome changes in other FASII components, leading to fitness improvement.

There is a notable number of populations containing a duplication of approximately 103 kb (coordinate: 22,738-125,834) in the *ΔfabH, ΔfabF,* and *ΔfabG* mutants, which includes one ORF encoding an acyltransferase and one ORF encoding acyl carrier protein. Additionally, a duplicated region of 10.1 kb (coordinate: 1,093,824-1,103,958) is shared by *ΔfabZ #1* and *ΔfabZ #3* and contains acyl-acyl carrier protein thioesterase. We hypothesize that these duplications may enhance fatty acid synthesis when *fabK, fabZ, fabH, fabF, or fabG* is defective.

#### ΔdpcK suppressors

DPCK is the final enzyme in the five-step pathway for CoA biosynthesis. ^49^ In the evolved *ΔdpcK* populations, mutations ≤100 bp upstream of *acpS* (encoding holo-ACP synthase) are present in eleven of the seventeen *ΔdpcK* populations. Holo-ACP synthase, responsible for covalent attachment of CoA to acyl carrier protein, uses CoA as a substrate for fatty acid biosynthesis. ^50^ Among the seventeen populations, one population, *ΔdpcK#17,* contained a gene duplication; we were not able to identify the gene(s) responsible for improved growth due to the large size of the duplication (>300 kb, containing 305 ORFs; Fig. 2i). We hypothesize that holo-ACP synthase is a significant contributor to the use of CoA, and when there is a shortage of CoA, fatty acid production needs to be reduced by lessening the expression of *acpS* (Fig. 3b and Supplementary Fig. 7, Supplemental Table 4 and Supplemental Table 15). Consistent with the fitness improvement of *<u>ΔdpcK</u>* by reducing CoA usage, seven out of the seventeen *ΔdpcK* populations contain mutations in the FASII operon (Fig. 3b and Supplementary Fig. 7 and Supplemental Table 15).

#### ΔpfkA suppressors

*pfkA* encodes 6-phosphofructokinase (PFK), responsible for catalyzing the phosphorylation of fructose-6-phosphate to fructose-1,6-bisphosphate, a committing step in glycolysis. ^51^ In the evolved *ΔpfkA* populations, we observed an enrichment of mutations in *fruR, fruB,* and *glck*. Among the ten *ΔpfkA* evolved populations (Fig. 3b, Supplementary Fig. 7, Supplemental Table 4 and 15), six populations contained mutations in *fruR*, five populations had mutations in *fruB*, and nine populations had mutations in *glcK*. The *glcK* gene encodes an ROK domain-containing glucokinase, which phosphorylates glucose to produce glucose-6-phosphate, a progenitor of fructose-6-phosphate. ^52^ The *fruR* gene encodes a transcriptional repressor of the *fruBA* operon, while *fruB* (also known as *pfkB*) has been demonstrated to function as a 1-phosphofructokinase in multiple bacterial species, ^53^ including streptococci. ^54^ Interestingly, the nature of the mutations differed according to the gene; three of the eleven *glcK* mutations were truncations, and four of the seven *fruR* mutations were truncations or frameshift mutations, while none of the seven *fruB* mutations were truncations or frameshifts (Supplemental Table 4 and Supplemental Table 15).

We hypothesize that in the absence of *pfkA*, accumulated fructose-6-phosphate becomes toxic, as has been shown previously in other bacterial species. ^53^ Elimination of GlcK activity would reduce fructose-6-phosphate production. We further hypothesize that FruB possesses low levels of fructose-6-phosphate kinase activity, thus replacing PfkA, but only when the expression of *fruB* is increased due to *fruR* mutation or the fructose-6-phosphate kinase activity of FruB is increased due to alteration of its sequence. This mechanism would be similar to findings in *E. coli*, where a mutation that increased *fruB* expression was found to suppress the phenotype of *ΔpfkA* mutants. ^55,56^

#### ΔobgE suppressors

ObgE belongs to the TRAFAC (Translation Factor Association) class of P-loop GTPases, which is highly conserved and essential in bacteria. ^57^ The ObgE GTPase is known to play a wide range of functions, with some of the most significant roles including sporulation initiation in *Bacillus subtilis*, ^58^ aerial mycelium formation in *Streptomyces*, ^59,60^ bacterial persistence in *E. coli* and *Pseudomonas aeruginosa*, ^61^ involvement in ribosome biogenesis, ^62,63^ participation in DNA replication, ^64–66^ and contributions to chromosome segregation and cell division. ^67^

Among the fourteen evolved *ΔobgE* populations, eight populations had substitution mutations upstream of *rpoB* (seven were an A to T mutation 321 bp upstream of the start codon, and the other was an A to G mutation 24 bp upstream), and two populations contained substitution mutations within the *rpoB* ORF (one at 905 bp and the other at 1058 bp from the start of the 3567 bp ORF). In total, 71.4% of the populations had mutations within or upstream of *rpoB* (Fig. 3b, Supplementary Fig. 7, Supplemental Table 4 and 15). The *rpoB* gene codes for the beta-subunit of RNA polymerase I. In one of these populations, *ΔobgE#5*, a 37 kb gene duplication was observed (coordinate: 1,775,394-1,812,805), encompassing thirty ORFs, including one annotated as Der GTPase. It was reported that overexpression of *der* or *obgE* can suppress growth defects of a mutant lacking *rrmJ* that encodes the methyltransferase for U2552 of the 23S rRNA. ^68^ From this, we hypothesize that the duplication may partially compensate for the loss of ObgE function by increasing *der* copy numbers, and a reduction in RNA synthesis is a significant fitness adaptation when ObgE is defective.

#### WT and other mutant suppressors

Suppressor mutations in the evolved populations of *ΔpngM*, *ΔyqeG, ΔdnaK, ΔyihA, ΔrpmH, ΔrplV,* and WT are less easily explained, and further study is required to identify possible mechanisms (Supplementary Fig. 7 and Supplemental Table 15).

### Testing the working model involving *f1fo, v1vo, trkA1-H1* and *trkA2-H2*

To test the hypotheses proposed above, we have chosen F1Fo-ATPase, an evolutionarily ancient enzyme complex that is highly conserved across virtually all forms of cellular life ^69^ and has been reported as essential for entire genus of the streptococcal species. ^35^

We began by testing whether increasing copy numbers of *v1vo* contributes to the fitness improvement of the mutant deleted for the entire F1Fo complex or single subunits. To explore this, we examined the gene duplication of the *v1vo* region at the early stage, P1 in the four populations that contain gene duplications at P6 (Fig. 2b-c, Supplementary Fig. 4b-c, and Supplementary Fig. 8a). Among these four populations, one had the gene duplication at P1, while the other three acquired the duplication during subsequent passages (Figure 4a and Supplementary Fig. 8a). This observation indicates that gene duplications of the *v1vo* region evolved in these populations after initial passages.

**Figure 4.**
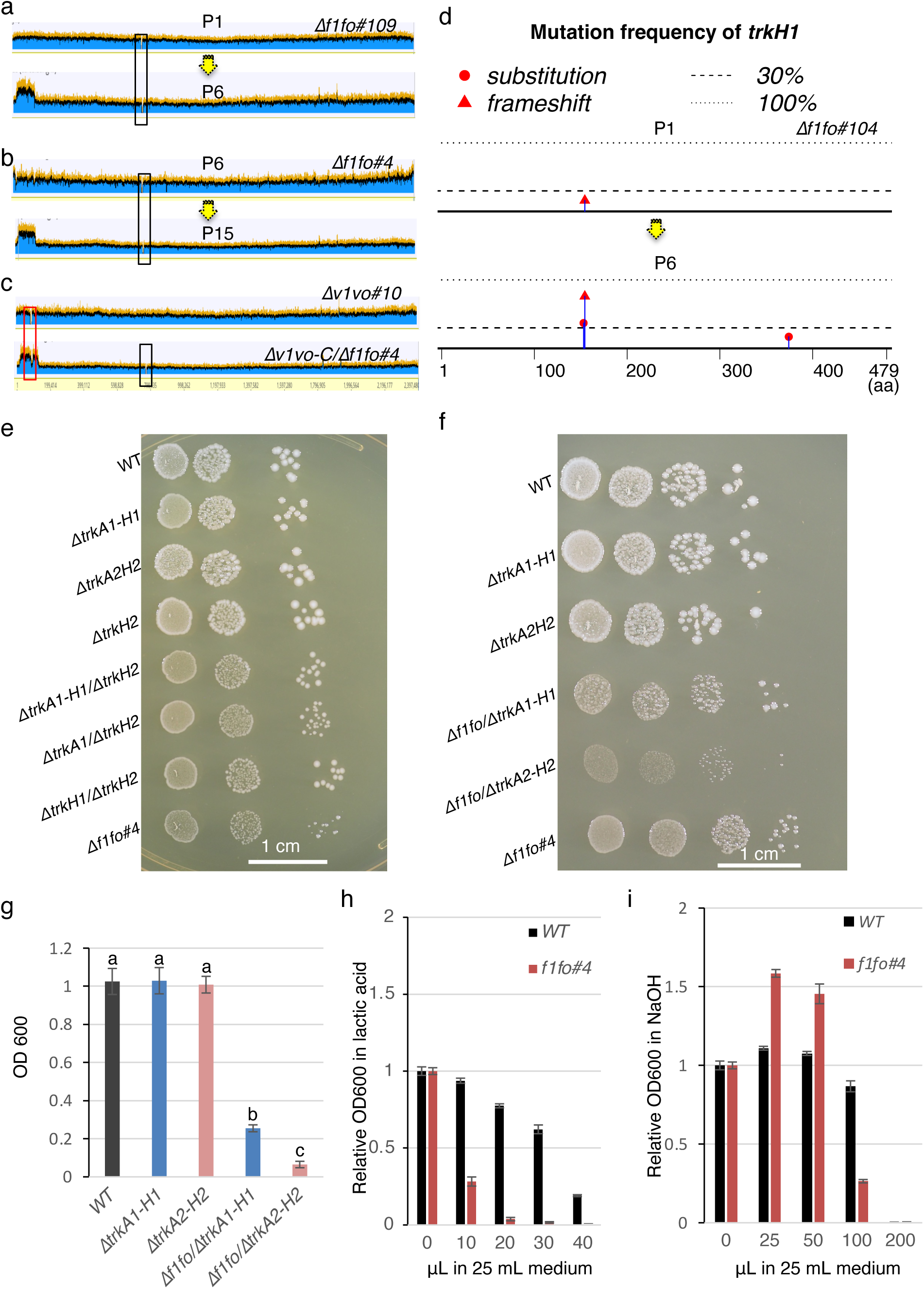
Increasing *v1vo* copy number and inhibition of *trkA1-H1* independently contribute to the fitness improvement of evolved *Δf1fo* populations. (a) Gene duplication of *v1vo* region does not occur in P1 population (upper) and appeared in P6 (lower) population of *Δf1fo#109.* Yellow arrows indicate passaging process. The black box indicates the *f1fo* region. (b) Population of *Δf1fo#4* at P6 that does not contain gene duplication of *v1vo* region (upper). *v1vo* region duplication was observed in P15 (lower). (Two out of sixed evolved populations; see S7). Arrows indicate passaging process. (c) *v1vo* region deleted in WT (upper). Duplication of *v1vo* region *Δf1fo#4, denoted as v1vo-DB/ Δf1fo#4.* The black box indicates the *f1fo* region. The red box indicates *v1vo* region. (d) Mutation frequency in *trkH1* increased with passage in P6 compared to P1 in population *Δf1f*#104. (e) Growth of strains indicated. 0.1 OD_600_ of cells in a volume of 2 µl were spotted directly (first column) or diluted 20-fold (column 2), or 400-fold (column 3) and grown on BHI-agar for two days. (f) Growth of the strains indicated. 0.1 OD_600_ of cells in a volume of 2 µl were spotted directly (first column) or diluted 10-fold (column 2), or 100-fold (column 3), or 1000-fold (column 4), and grown for four days. (g) Quantitative growth measurements of WT, *ΔtrkA1-H1, ΔtrkA2-H2, Δf1fo* in *ΔtrkA1-H1* and *Δf1fo* in *ΔtrkA2-H2.* Culture is in BHI for 24 hr in anaerobic conditions. Data are shown as means ± SD from three replicate cultures. Different letters indicate statistically significant differences (P≤0.05), determined by one -way ANOVA with Tukey multiple comparisons test. (h) Sensitivity of *Δf1fo#4* and WT to low pH (lactic acid: LA). The values are relative OD_600_ to that of WT grown in plain BHI. X-axis denotes the vol (μL) of 90% LA in 25 mL of BHI. Data are shown as means ± SD from three replicate cultures. (i) Sensitivity of *Δf1fo#4* and WT to high pH (NaOH). The values are relative OD_600_ to that of WT grown in plain BHI. X-axis denotes the vol (μL) of 5 M NaOH in 25 mL of BHI. Data are shown as means ± SD from three replicate cultures.

To further confirm the increased fitness of *v1vo* region duplications in populations lacking *f1fo*, we subcultured five *Δf1fo#4* populations from the P6 stage, which did not initially contain gene duplications in the *v1vo* region. After nine additional passages (P15), we observed that two out of the five evolved populations had *v1vo* region duplications in P15 (Fig. 4b and Supplementary Fig. 8a).

To ascertain the fitness improvement by increasing *v1vo* copy numbers in mutants lacking *f1fo*, we tested whether *v1vo* could be deleted in *Δf1fo*. We first deleted the entire *v1vo*, which consists of nine subunits, to generate the *Δv1vo* mutant in the WT background. This *Δv1vo* mutant grew indistinguishably from the wild type in a rich medium (data not shown), demonstrating that *v1vo* can be deleted when *f1fo* is intact (Fig. 4c). However, when we attempted to delete *v1vo* in *Δf1fo#4*, where *f1fo* was removed, it resulted in a double-band genotype in all five *Δv1vo-DB/Δf1fo#4* ("DB" for double-band) lines tested (Fig. 4c and Supplementary Fig. 9). Consistent with these findings, *ΔuncE#2* and *ΔuncH#7*, which contain amino acid substitution mutations of V1Vo ATP synthase subunit A and K in the *v1vo* operon, respectively, also contain *v1vo* region duplications (Supplemental Table 4). This finding indicates that increased copy numbers of the *v1vo* genes compensated for the loss of F1Fo function. Overall, these results suggest that increasing the copy numbers of *v1vo* contributes to the fitness improvement of mutants lacking *f1fo*.

We then investigated the impact of inhibiting *trkA1-H1* on enhancing the fitness of *Δf1fo*. Initially, we compared the occurrence of *trkA1-H1* mutations between P1 and P6 stages across twelve independently evolved populations of *Δf1fo* (*#101* to *#112*). While mutations in the *trkA1* or *trkH1* segments were not detected in P1 using a 30% threshold in nine of the twelve populations, we observed *trkA1-H1* mutations in all populations, with the mutation frequency increasing during passage in all six independently evolved populations (Fig. 4d and Table S17). These mutations resulted in at least one frameshift and various substitutions in the translation of TrkA1 or TrkH1 in six populations, while in the remaining populations, they led to a histidine insertion or various substitutions in the TrkH1 protein (Supplemental Table 16). This data indicates the fixation of *trkA1-H1* mutations during passage.

To further investigate the impact of *trkA1-H1*, we compared the growth of *Δf1fo* in the presence and absence of *trkA1-H1*. *S. sanguinis* SK36 has two Trk systems, *trkA1-H1* and *trkA2-H2*. The growth of *trkH2*, or double-gene deletion mutants of *trkA1-H1* or *trkA2-H2,* were indistinguishable from that of WT (Fig. 4e, Supplementary Fig. 10a-f and Supplemental Table 17). However, combined deletion of either *trkA1* or *trkH1* along with *trkH2* resulted in growth defects (Fig. 5e-g, Supplementary Fig. 10g-h, and Supplemental Table 17), indicating that *trkA1-H1* and *trkA2-H2* are redundant for growth. In order to determine the effect of mutating the two Trk systems on the fitness of *Δf1fo*, we transformed the *Δf1fo* construct into the backgrounds of *ΔtrkA1-H1*or *ΔtrkA2-H2*, and compared the growth of the *Δf1fo/ ΔtrkA1-H1* mutant and that of *Δf1fo/ΔtrkA2-H2*. As controls, we also included *Δf1fo#4*, which contains a frameshift mutation of *ΔtrkH1* in the *Δf1fo* background (Supplemental Table 4). It showed that while the growth of *Δf1fo#4* and *Δf1fo/ΔtrkA1-H1* were comparable to each other, the growth of *Δf1fo/ΔtrkA2-H2* was significantly inhibited compared to that of *Δf1fo#4* and *Δf1fo/ΔtrkA1-H1.* (Fig. 4f-g and Supplemental Table18). These findings suggest that inhibiting *trkA1-H1*, but not *trkA2-H2*, contributes to the fitness improvement of evolved mutants deleted for *f1fo*.

**Figure 5.**
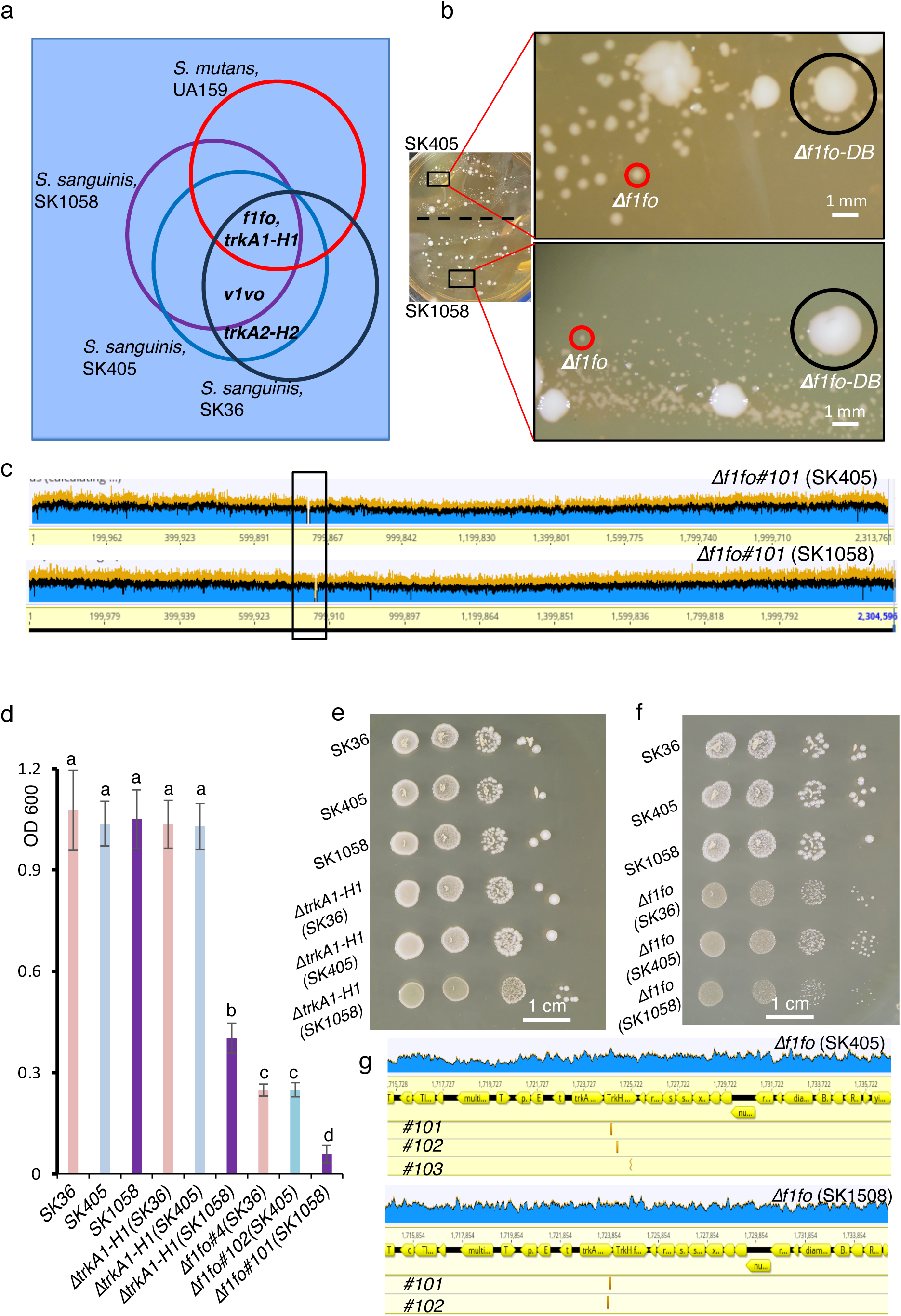
Essentiality of *f1fo* in *S. mutans* UA159, *S. sanguinis* SK36, SK405 and SK1058. (a) Distribution of the genes indicated in *S. mutans* UA159, *S. sanguinis* SK36, SK405 and SK1058. (b) Growth of *Δf1fo* transformants in *S. sanguinis* SK405 (upper) and SK1058 (lower) on same BHI agar medium for five days. Enlarged images of the transformants are shown. Red circles indicate small colonies, *Δf1fo*. Black circles indicate big colonies, *Δf1fo-DB*. (c) Deletion of *f1fo* in SK405 or SK1058 shown by whole genome sequencing. Black box indicates the location of *f1fo* region. (d) Quantitative measurements of growth in wild-type SK36, SK405 or SK1058, *ΔtrkA1-H1* or *Δf1fo* mutants in the backgrounds of SK36, SK405 or SK1058. Cells were grown in BHI for 24 hours. Data are shown as means ± SD from three replicate cultures. Different letters indicate statistically significant differences (P≤0.05), determined by one -way ANOVA with Tukey multiple comparisons test. (e) Growth of SK36, SK405 or SK1058, *ΔtrkA1-H1* mutant in the backgrounds of SK36, SK405 or SK1058. 0.1 OD_600_ of cells in a volume of 2 µl were spotted directly (first column) or diluted 10-fold (column 2), or 100-fold (column 3), or 1000-fold (column 4). Growth of Cells were grown on BHI-agar for two days. (f) Growth of SK36, SK405 or SK1058, *Δf1fo* mutant in the backgrounds of SK36, SK405 or SK1058. Cells were diluted as in Fig. E and grown on BHI-agar for four days. (g) Location of *trkA1-H1* suppressor mutations in SK405 and SK1058. Mutation of *trkA1-H1* appeared in all evolved *Δf1fo* populations, three in SK405 and two in SK1058. X-axis indicates the genome coordinates.

In lactic acid bacteria, F1Fo-ATPase is believed to function by pumping out protons to alkalinize the cytosol or to build negative internal and positive external membrane potentials. ^70^ To test whether the growth defects in the *S. sanguinis Δf1fo* mutant are due to impaired function in alkalinizing the cytosol or building a membrane potential, we compared the growth of *Δf1fo#4* in a rich medium supplemented with lactic acid (which increases membrane potential and cytosolic proton concentration) or NaOH (which lowers membrane potential and cytosolic proton concentration). We observed that *Δf1fo#4* is susceptible to lactic acid treatment compared to WT (Fig. 4h and Supplemental Table 18). Conversely, the growth defect of *Δf1fo#4* can be partially rescued by NaOH within a certain range (Fig. 4i and Supplemental Table 18). These results suggest that *f1fo* in *S. sanguinis* functions by pumping out protons to alkalinize the cytosol.

We therefore suggest the following model for the relationship of *f1fo* with *v1vo, trkA1-H1,* and *trkA2-H2* (Supplementary Fig. 11). For streptococci, which do not contain the tricarboxylic acid cycle or electron transport chain, energy production is mainly through lactic acid fermentation. ^71^ The lactate from fermentation creates highly acidic cytosol, which would be toxic if H+ were not exported. F1Fo pumps out H+ from the cytosol using ATP. Both TrkA1-H1 and TrkA2-H2 are potassium transporters and functionally redundant for that purpose. TrkA1-H1 pumps in potassium together with H+, acidifying the cytosol when H+ is not exported. In contrast, TrkA2-H2 does not pump protons into the cytosol when pumping in potassium (likely instead pumping in Na+). Therefore, the function of TrkA1-H1 must be inhibited if F1Fo is not functional.

### Conserved functional relationships of *f1fo*, *trkA1-H1, trkA2-H2* and *v1vo* in different streptococci

To generalize the *f1fo* working model and assess F1Fo’s essentiality in other strains and species, we conducted essentiality evaluations in three other streptococci: *S. mutans* UA159, *S. sanguinis* SK405, and *S. sanguinis* SK1058. Genomic sequencing of *S. mutans* UA159, ^72^ *S. sanguinis* SK405 ^38^, and *S. sanguinis* SK1058 ^38^ showed that *f1fo* and *trkA1-H1* are present in all three strains, as in SK36, while *v1vo* is present in the *S. sanguinis* strains only. *TrkA2-H2* is present in both *S. sanguinis* SK36 and SK405 but not in *S. sanguinis* SK1058 or *S. mutans* UA159 (Fig. 5a).

To assess whether *v1vo* could serve as a partial backup in cases of F1Fo loss-of-function, we attempted to delete *f1fo* in *S. mutans* UA159, which lacks *v1vo*. Despite multiple attempts and a total of 64 independent genotyping analyses, none of the transformants were deleted for *f1fo*, instead producing a double band indicative of gene duplication prior to allelic exchange (Supplementary Fig. 12a-b). This result highlights that in *S. mutans* UA159, lacking *v1vo, f1fo* cannot be deleted in our conditions. This observation supports the evolutionarily conserved relationship between *f1fo* and *v1vo* observed in other streptococcal species.

We then investigated the impact of deleting *trkA1-H1* in the other two strains: *S. sanguinis* SK1058, which only possesses *trkA1-H1*, and *S. sanguinis* SK405, which has both *trkA1-H1* and *trkA2-H2*. Our findings showed that *trkA1-H1* mutants exhibited severe growth defects in the SK1058 background compared to the SK405 background (Fig. 5d-e and Supplementary Fig. 13a-c). This result demonstrates the requirements of the *Trk* system for growth within the species of *S. sanguinis*.

Next, we investigated the possibility of deleting *f1fo* in *S. sanguinis* SK1058 and SK405, both of which contain *v1vo* and *trkA1-H1*. We could generate *f1fo* whole-region deletion mutants (*Δf1fo*) in both SK1058 and SK405 backgrounds (Fig. 5b, 5c, Supplementary Fig. 14). Furthermore, the growth of mutants with *f1fo* deletions in SK405 was notably better than in SK1058 (Fig. 5b, 5d-f and Supplemental Table 19). To explore the potential emergence of *trkA1-H1* suppressor mutations during evolution, we conducted passage experiments using independently evolved populations with *f1fo* deletions in SK405 (Supplementary Fig. 14a: three populations) and in SK1058 (Supplementary Fig. 14b: two populations). It was observed that, after six passages, *trkA1-H1* suppressor mutations appeared in all evolved *Δf1fo* populations in SK405 and SK1058 (Fig. 5e). At least one population each in the SK405 and SK1058 backgrounds are frameshift or truncation mutations of *trkA1-H1* (Fig. 5g and Supplemental Table 20). This result underscores the evolutionarily conserved occurrence of *trkA1-H1* suppressor mutations in evolved populations with *f1fo* deletions.

## Discussion

While identifying genes that interact with essential genes or compensate for their loss provides a great benefit, ^4^ this task remains challenging, primarily due to the unavailability of essential-gene deletion mutants and systematic methodologies to connect the function of essential genes with their partners. Traditional approaches, such as conditional promoter-shutoff^73,74^, antisense RNA knockdowns ^48^ and CRISPR interference, ^75,76^ while valuable, may exhibit limitations, including incomplete target gene silencing and suboptimal design efficacy, ^76^ thereby potentially compromising the reliability of essential gene functional analyses. Our study successfully established a system to generate viable mutants with deletions in essential genes (Fig. 1 and Supplementary Fig. 1-S2). We observed that small colonies likely represent the deletion mutants, while large colonies often exhibit the "double-band" genotype or kanamycin resistance due to mutations in translational machinery.

With these observations, we have generated essential gene deletion mutants, many of which are also essential across multiple species (http://www.essentialgene.org/). These essential gene mutants provide an invaluable resource for dissecting the functions of essential genes. Using these twenty-four essential gene mutants and WT as controls, we identified over 1000 spontaneous suppressor mutations in passage experiments (Fig. 2). Leveraging these essential gene mutants and distinct mutated segments, we constructed a functional network (Fig. 3a). Within this network, we made several noteworthy observations. 1) Mutated segments that appeared in the evolved WT population can also appear in the evolved populations of mutants with deletions of essential genes, such as mutations in the PTS mannose/fructose/sorbose transporter family subunit. 2) Suppressor mutations in certain genes, such as *rpoB* and *rpsL,* that are found in the evolved populations of one deletion mutant can also be present in the evolved populations of other mutants (Supplemental Table 4). 3) Mutual essential gene suppressor mutations were discovered, where mutations in the first essential gene were found within a mutant population deleted for the second essential gene, and vice versa.

As expected, our findings support the notion that mutants with deletions of essential genes face stronger selective pressures when compared to that of the WT. These mutants are more likely to undergo the following significant evolutionary changes. 1) Gene duplications of greater than 10 kb: Our data revealed large-scale gene duplications exclusively were present in mutants deleted of essential genes (Fig. 2b-h). 2) Substitutions, insertions, and deletions of less than 10 kb: Small-scale mutations, especially those of >50% of sequence reads, were more common in mutants deleted for essential genes (Fig. 2i). 3) Mutations in essential genes: Mutations in other essential genes were observed more exclusively in mutants deleted for an essential gene (Fig. 2k). 4) Mutations in multi-drug resistance genes: mutations of multi-drug resistance-associated genes, such as *rpoB* and *rpsL,* are present in several evolved populations deleted for essential genes, suggesting a shared selective pressure resulting from drug treatment and deletion of essential genes (Supplemental Table 4). The suppressor mutations identified in our evolved populations deleted of essential genes may reflect the differential gene essentiality over evolutionary time in the pan-genome ^11^ and reproducibly changing gene essentiality across independently evolving populations. ^18^

Unexpectedly, as discussed above, for every essential gene mutated, we identified at least one population containing a suppressor mutation in another essential gene (Fig. 2l). Notably, the essential genes were inactivated by the suppressor mutation in some of the evolved populations, such as the >200bp insertion mutation in *ftsZ* in the evolved population of *Δf1fo #107*, the >200 bp deletion mutation of *uncH* in *ΔuncC#1*, and the 1 bp deletion mutation of *ftsE* in *ΔftsX#13* (Supplemental Table 4). In contrast, for other essential gene mutation suppressors, the mutations were not within the open reading frame (ORF) but were found within 400 base pairs upstream of the start codons, likely affecting expression rather than inactivating function. For instance, in the evolved *ΔdpcK* populations, the suppressors with the highest mutation frequency were located upstream of the holo-ACP synthase gene, *acpS* (Supplementary Fig. 5). Similarly, in the evolved *ΔobgE* populations, the majority of mutation suppressors were found upstream of *rpoB* (Supplementary Fig. 5).

Essential genes represent ideal drug targets for controlling infectious diseases and cancers. In traditional medical treatment, the focus has primarily been on targeting a single essential gene product. However, it’s crucial to consider the evolution of gene essentiality during drug treatment and the development of drug resistance over short-term evolution. We have observed that in multiple populations deleted of essential genes, mutations in multiple segments appeared within six passages, including those belonging to multi-drug resistance-associated genes (Fig. 2m). Considering that some of the suppressor mutations were gene duplications that presumably confer fitness benefits by increasing gene copy numbers, a promising approach is to target not only the essential gene itself but also the backup gene. This strategy can help prevent or reduce the development of drug resistance.

To understand the relationships between the structure and function of a protein, it is of great interest to know which amino acids or domains of the proteins are sensitive to mutations. In this study, we have observed numerous mutations of *trkA1* and *trkH1* in *Δf1fo* or *f1fo* single subunit deletion mutants, *vicK* in *ΔftsX*, *fruR* and *glcK* in *ΔpfkA*, as well as *fusA*, *rpsL* and *rplF* in kanamycin-resistant mutants lacking the *kan* gene (Supplemental Table 1-2). These mutations represent fitness-related protein sequence changes that improve the growth of resultant mutants in a particular selective condition.

In this work, we have demonstrated the efficacy of the methodology developed for generating deletion mutants of numerous essential genes in *S. sanguinis* SK36. Furthermore, we have highlighted the potency of experimental evolution ^29,30^ in unraveling the mechanisms of gene essentiality, with a particular focus on *f1fo* in *S. sanguinis* and *S. mutans*. It would be interesting to test the conservation or divergence of mechanisms governing gene essentiality of other genes, as well as across different *Streptococcus* species and other genera. While extracting valuable information by identifying spontaneous suppressor mutations in populations lacking essential genes is possible, conducting experimental evolution requires caution. First and foremost, the choice of an appropriate organism is crucial. Ideally, the selected organism should have a single chromosome, a small genome size, and a rapid cell division rate. Streptococci represent an ideal candidate in this regard. They possess a single circular genome with a relatively modest size, typically ranging from 2.0 to 3.0 megabases. ^77^ This choice contrasts with a previous study that employed budding yeast with multiple chromosomes ^4^ where most mutations in populations lacking essential genes were related to changes in ploidy, which could hinder the identification of causal suppressors. On the other hand, the identification of mechanisms of genetic suppression requires caution. For the suppressor mutations of low frequency or these substitution mutations where we were unable to determine whether the evolved suppressor mutations increase or decrease the activity of the gene product, further evidence is needed to understand the mechanisms underlying the evolution of gene essentiality.

In general, essential genes tend to be more conserved and evolutionarily ancient compared to non-essential genes. ^54^ It is intriguing to consider when genes emerge within a biological system over time. The suppressor mutations in essential gene mutants, particularly the loss-of-function mutations, can provide valuable insights into dependency relationships. For instance, the absence of *f1fo* in a system may hinder the emergence of the *trkA1-H1* system. In cases where essential genes function as a complex, such as *f1fo* or *ftsEX*, our findings indicate that deletion of one component may result in additional deleterious effects caused by the remaining subunit(s) (Supplemental Table 4). This suggests that all subunits of the complex need to be integrated into the system, at least for these cases. Moreover, understanding the timelines of gene emergence in a system is also crucial for guiding decisions in synthetic biology, determining which genes, as well as the order of genes, to be introduced into a system.

## Materials and Methods

### Strains Used and Growth Conditions

In this study, we primarily utilized *Streptococcus sanguinis* strain SK36. ^32^ *S. sanguinis* SK405 and SK1058 strains were described previously. ^38^ *S. mutans* UA159 was obtained from Ann Progulske-Fox (University of Florida). All bacterial cultures were routinely maintained in a BHI medium at 37°C under anaerobic conditions, using a Coy anaerobic chamber, or stored at -80 °C for long-term storage.

### Transformation and Mutant Isolation

To prepare competent cells, *S. sanguinis* was cultured in 1 mL of Todd-Hewitt + HS medium ^10^ and incubated overnight in anaerobic conditions at 37°C. The following day, a 5 μL aliquot of the overnight culture was added to 1 mL of fresh TS+HS medium and incubated for an additional 3 hours.

For creating knockout constructs, we employed 1 kb of homologous 3’ and 5’ end sequences flanking the coding sequence (CDS) and resistance genes, *kan* or *Erm*^r^, in overlapping extension PCR (for primers used, see Supplemental Table 21). ^32^ For transformation, we mixed 50 ng to 500 ng of PCR product DNA with 200 ng of CSP and 300 μL of competent cells. The reactions continued for 24 hours under anaerobic conditions at 37°C. Subsequently, transformation reaction mixtures were plated on BHI-agar containing appropriate antibiotics (Kanamycin: 500 μg/mL or Erythromycin: 10 μg/mL) for selection. Once the plates had dried, the cells were overlaid with a thin layer of agar by adding 1 mL of cooled BHI-agar with antibiotics. The plates were incubated at 37°C under anaerobic conditions to select transformants for 4 to 5 days.

For genotyping, genomic DNA (gDNA) was isolated from 2 mL of saturated cultures. In brief, cells were precipitated by centrifugation at 10,000 rpm for 10 minutes at room temperature. They were then resuspended in 200 μL of resuspension buffer (20 mM EDTA, 200 mM Tris-HCl, and 2% Triton-X 100) and lysed with 200 μL of AL lysis buffer (Qiagen, 19075). DNA was precipitated by adding 1 mL of 100% ethanol supplemented with 100 mM of sodium acetate. Following washing and drying, the DNA was dissolved in 100 μL of water and used for PCR and/or whole-genome sequencing.

To generate deletion mutants *Δf1fo* in *ΔtrkA1-H1* and *ΔtrkA2-H2*, the *ΔtrkA1-H1* and *ΔtrkA2-H2* mutants were first generated using *Erm*^r^ (Supplementary Fig.10). After confirmation of deletion of *ΔtrkA1-H1* and *ΔtrkA2-H2, Δf1fo* of *kan* resistance construct was subsequently transformed into *ΔtrkA1-H1* or *ΔtrkA2-H2.* To generate deletion mutants of both *trkA1-H1* and *trkA2-H2*, *trkA1, trkH1*, or *ΔtrkA1-H1* of erythromycin resistance constructs were transformed into a *ΔtrkH2* mutant of kan resistance background (Xu, et al., 2011). To isolate *Δf1fo* and *ΔtrkA1-H1* mutants in the SK405 and SK1058 strains, we employed the same PCR construct used for deleting *f1fo* and *trkA1-H1* in SK36, as there is >90% identity at the flanking regions between SK36, SK1058, and SK405.

### Strain Passage

Antibiotic-resistant colonies from selection plates were transferred to 1 mL of BHI containing antibiotics and grown for 2 to 8 days under anaerobic conditions at 37°C. The duration depended on when the OD_600_ reached 0.1-0.5, which was referred to as P0. A 300 μL aliquot of the cell culture was then transferred to 3 mL of BHI containing antibiotics, and growth continued until saturation, resulting in P1. P1 was mixed by pipetting five times with a P-1000 pipette, and 50 μL of the mixed P1 was transferred to 1 mL of BHI containing antibiotics. Growth proceeded until the cell culture reached an OD_600_ of 0.1 to 0.5, producing P2. The latter process was repeated sequentially from P3 to P6, where the volume of P2 to P5 was 1 mL and the final passage, P6 was 3 mL. Cell cultures of P1 and P6 were stored at -80°C in glycerol at the final concentration of 20%. DNA extractions and genome sequencing were conducted in P6 and/or P1 when indicated.

For passages involving WT, glycerol stocks from -80°C were transferred to 1 mL of BHI and grown for 24 hours, resulting in T0 (see Supplementary Fig.4a). A 50 μL aliquot of T0 cell culture was aliquoted into six 1.5 mL tubes, each containing 1 mL of BHI, and grown for 24 hours, resulting in T1. A 50 μL aliquot of T1 cell culture was inoculated into 1 mL of BHI and grown for 24 hours, leading to T2. This process was repeated for T3 to T9.

### Whole-Genome Sequencing and Variant Calling

Whole-genome sequencing was carried out by the Genome Core at Virginia Commonwealth University or by SeqCenter (https://www.seqcenter.com/) using the shotgun method with 2×150 paired-end sequencing. Whole-genome sequences with an average coverage of ≥100 were used for subsequent analysis.

Fastq files were aligned to the SK36 reference genome ^16^ using Geneious Prime software (https://www.geneious.com/) after trimming via the BBDuk method. Variations in the genome were exported from Geneious Prime. To identify the mutated segment, the frequencies (percentages) of all mutations belonging to a certain segment were assessed and the greatest value of percentages were assigned to that segment. For the deletions, the read coverages were calculated by averaging the base coverages that were 100 bp from the flanking sequence. For network analysis, the software Cytoscape was used.^78^

## Supporting information

Data.files.Experimental Evolution in Bacteria

## Data and materials availability

All data needed to evaluate the conclusions in the paper are present in the paper and/or the Supplementary files. The genome sequence data will be deposited to BioProject (PRJNA1146876). All the transgenic materials, including the essential gene deletion mutants are available from the authors upon request.

## Contributions

L.B. and P.X. conceived the project. L.B., T.K. and P.X. designed the experiments. L.B. constructed all the essential gene mutants and performed all experimental evolution studies. L.B., T.K., and P.X. analyzed the data with the help of Z.Z., A.I., B.Z., V.A. and M.W. The manuscript was written and revised by L.B., T.K., and P.X.

## Acknowledgments

The authors would like to thank Dr. Mark Running at the Department of Biology, University of Louisville, Dr. Tomomichi Fujita at the Department of Biological Sciences, Hokkaido University, Dr. Tim van Opijnen at Broad Institute, MIT and Harvard, and Dr. Yan Bao at the School of Agriculture and Biology, Shanghai Jiao Tong University, for discussions regarding experimental design and critical reading of the manuscript. We also thank Dr. Junling Ren at the Philips Institute for Oral Health Research, Virginia Commonwealth University for help with network analysis and Ms. Ayana Wyche at Virginia Commonwealth University for assistance with the culturing of bacteria. We are grateful to Vladimir Lee and Kalyan Maliempati for genomic sequencing services. This work was funded by grants from the National Institutes of Health R01DE030121 (P.X. and T.K.) and the CCTR Endowment Fund (P.X.).

## Declaration of generative AI in scientific writing

During the preparation of this work the authors used ChatGPT-3 in order to improve readability and language. After using it, the authors reviewed and edited the content as needed and took full responsibility for the content of the publication.

## Supplemental Figures, Tables and legends

**Supplementary Fig. 1.**
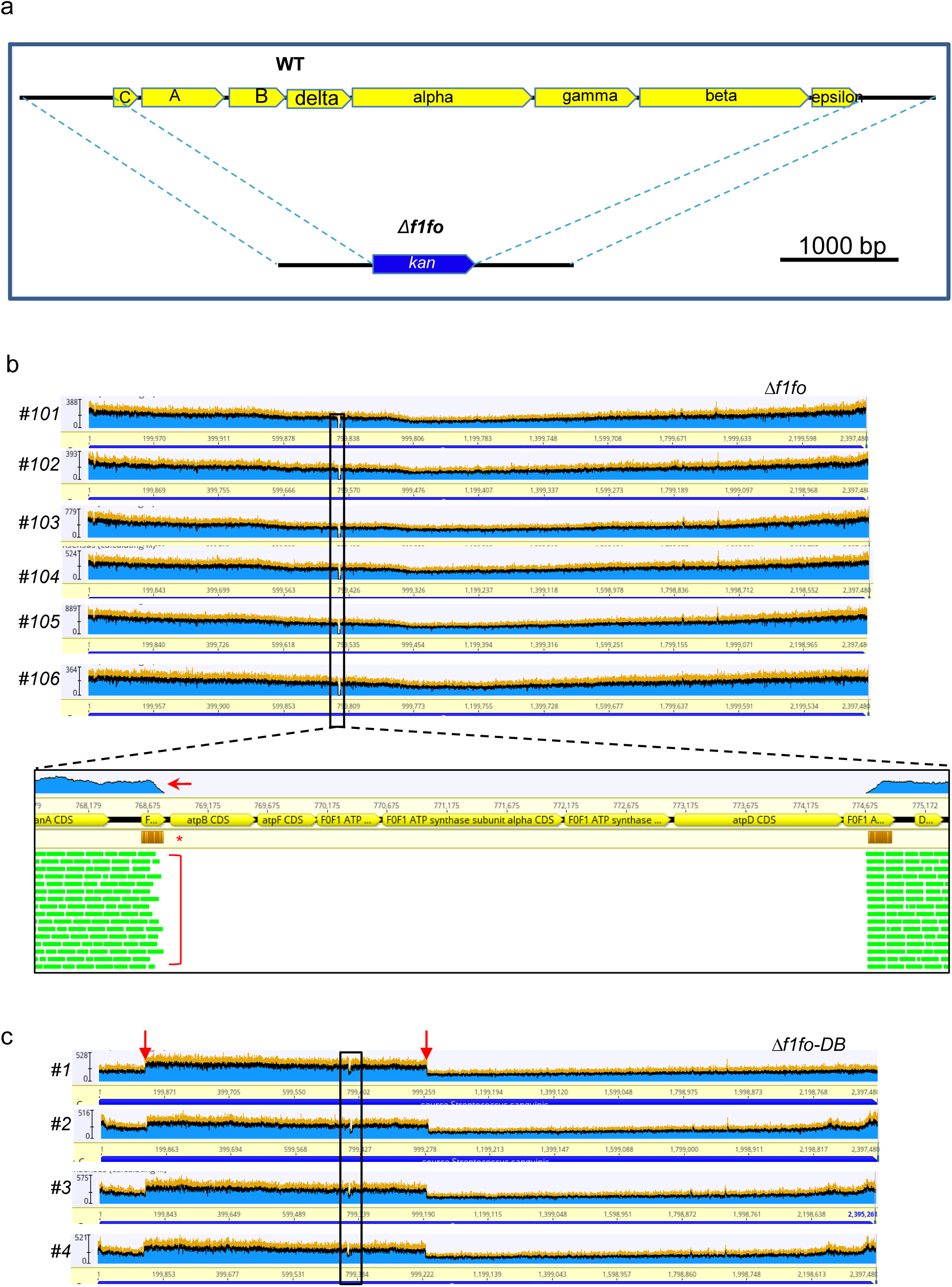
Genotyping and complementation of mutants deleted of *f1fo*. (a) Gene knockout strategy to delete entire region of eight *f1fo* subunits by homologous recombination. The yellow blocks indicate the eight F1Fo subunit genes in WT (upper). The blue block indicates the *kan* gene in *Δf1fo* mutants (lower). (b) Whole-genome sequencing to confirm the genotypes of the small colonies for *Δf1fo* mutants. Black box indicates the location of deleted *f1fo* region. Enlarged image showed alignment of sequencing reads to the reference genome of *S. sanguinis* SK36. The blue graphic above the sequence coordinates (indicated by a red arrow) indicates the extent of coverage, the yellow bars indicate the mismatches (indicated by a star), and the bracket indicates the aligned sequence reads. (c) Whole-genome sequencing of the big colonies, *Δf1fo*-DB. Black box indicates the location of *f1fo* region. Red arrows indicate the two directly repeated *ugpC* genes, encoding glycerol-3-phosphate ABC transporter.

**Supplementary Fig. 2.**
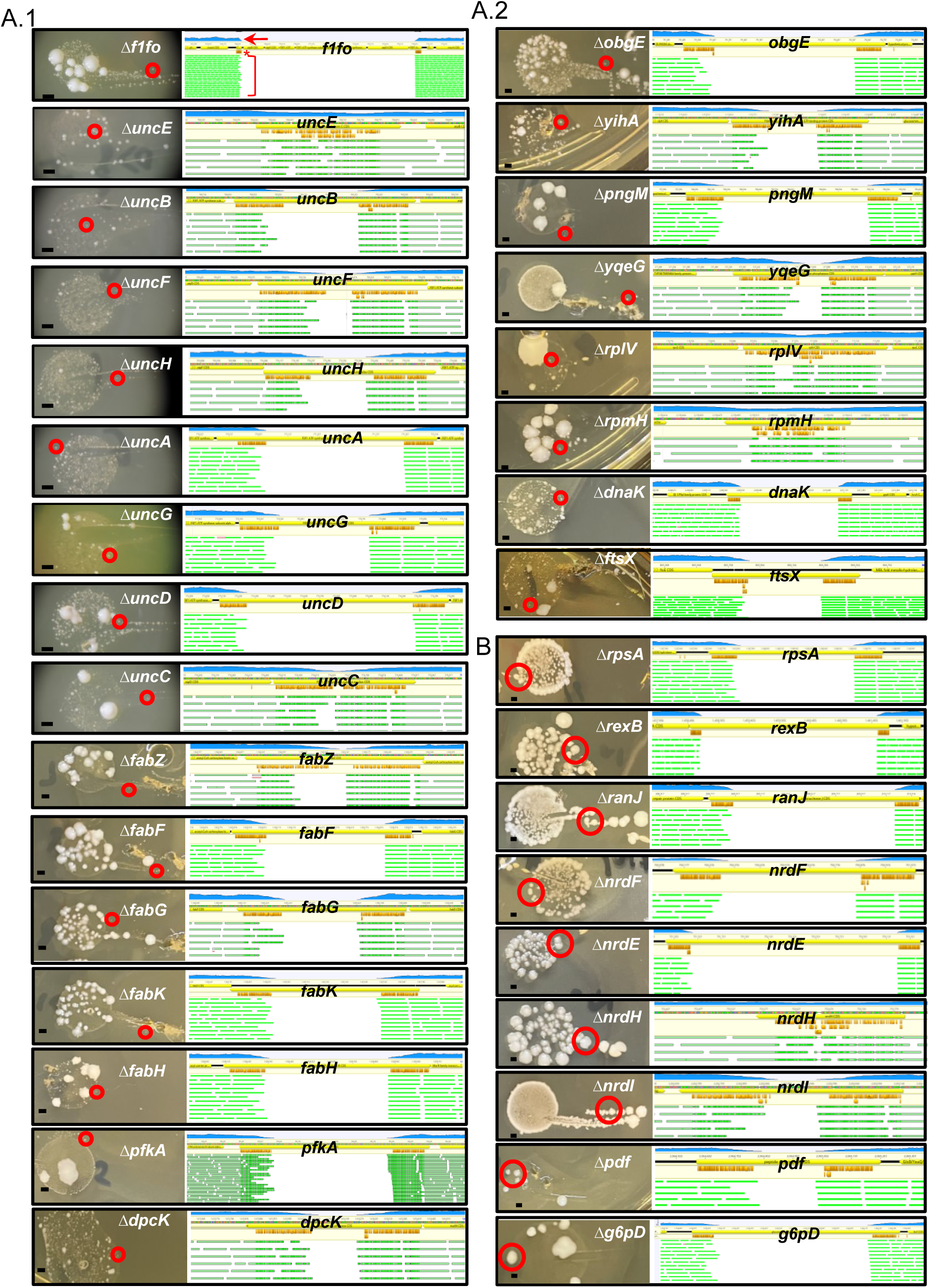
Isolation of viable mutants deleted essential genes and genotyping by whole-genome sequencing. (a-b) Colonies of transformants (left) and whole-genome sequencing of colonies (right). Red circles indicate the colonies picked for whole-genome sequencing (left). For genome sequencing (right), the blue graphic above the sequence coordinates (indicated by a red arrow in the top panel of a.1) indicates the extent of coverage, the yellow bars indicate the mismatches, and the bracket indicates the aligned sequence reads. Bacterial colonies on selective agar plate for five days. (a.1-a.2) Gene deletion mutants with severe growth defects. (b) Gene deletion mutants with robust growth. Scale bar: 1 mm. The image of *Δf1fo* transformation is a duplicate of Figure 1B to demonstrate the transformants in selection medium *Δf1fo* alignment is a duplication of Fig.S1c.

**Supplementary Fig. 3.**
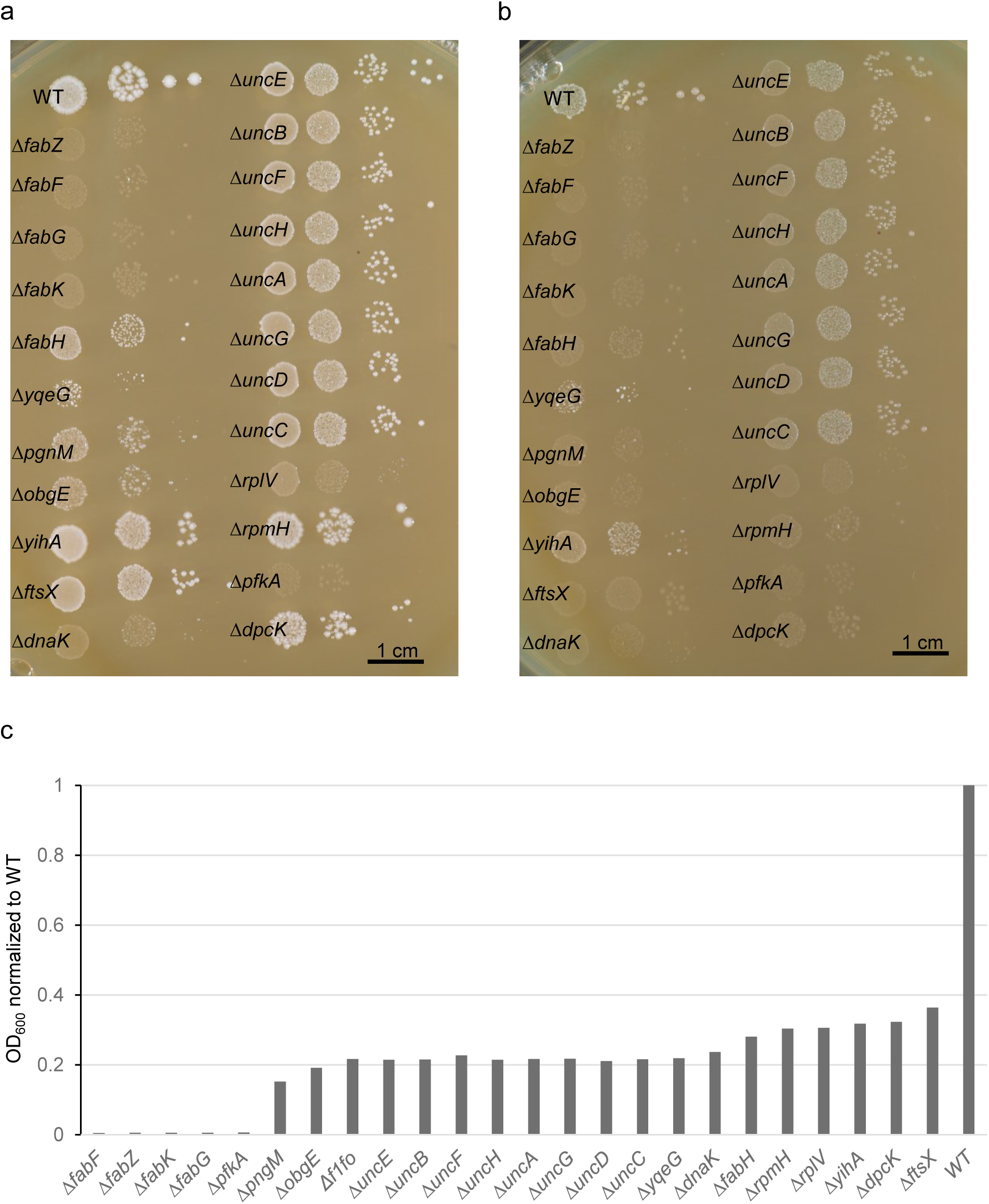
Growth of mutants deleted of essential genes, relative to that of WT. (a-b) Growth mutants deleted of 23 essential genes under anaerobic condition (a) and microaerobic condition (b). 0.1 OD_600_ of cells in a volume of 2 µl were spotted directly (first column) or diluted 20-fold (column 2), or 400-fold (column 3) and grown on BHI-agar for two days. Scale bar is 1 cm. (c) Comparison of the growth of the twenty-four evolved mutants and that of WT. The values are relative OD_600_ after 24 hr of growth in BHI liquid medium.

**Supplementary Fig. 4.**
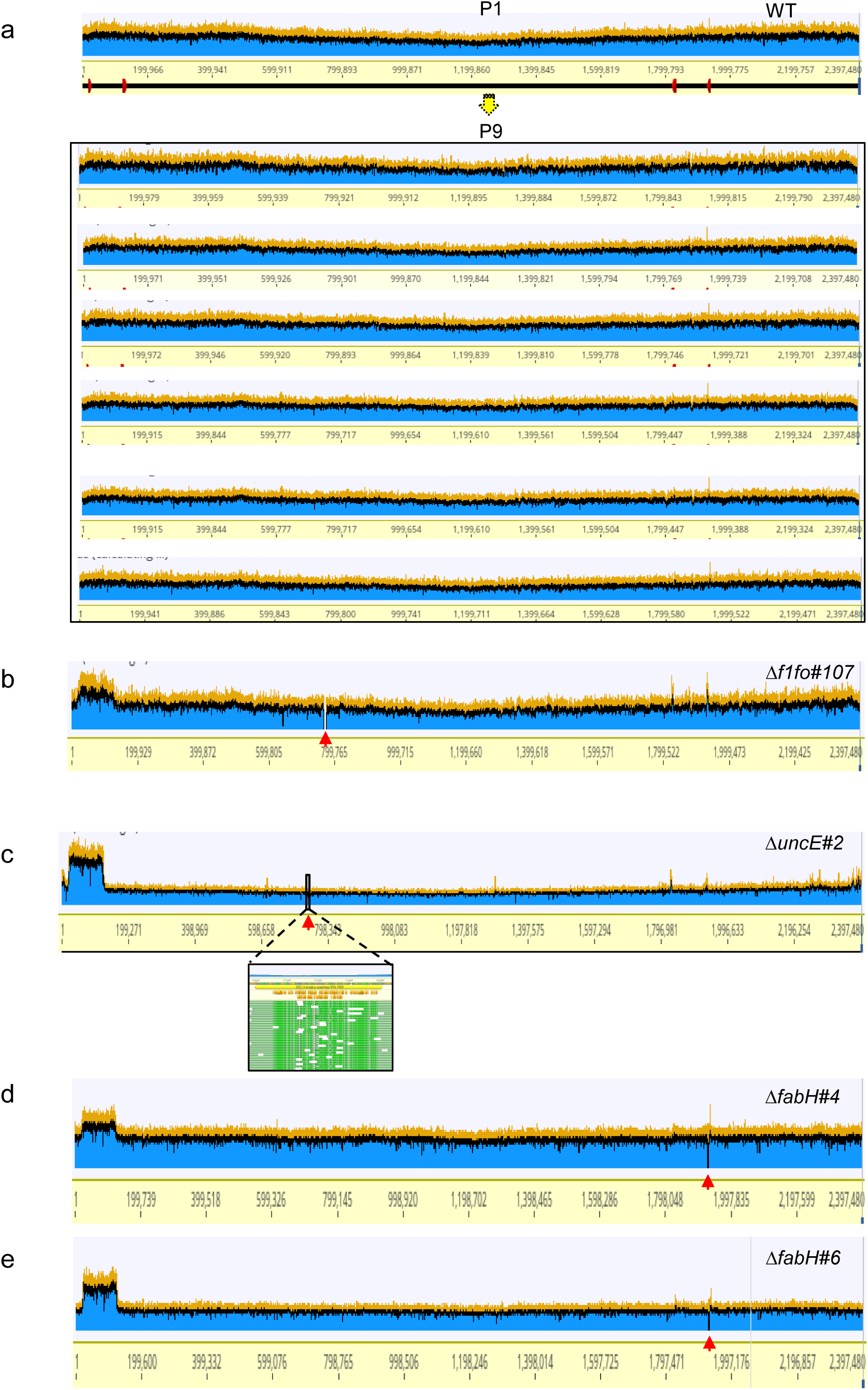
Large scale gene duplication present in the evolved populations of the mutants deleted essential genes but not in WT populations. (a) Whole-genome sequencing of parental WT and the six evolved WT populations after nine passages. The yellow arrow indicates the passage procedure. (b-e) Evolved populations that contain gene duplication of greater than 10 kb in *ΔuncE#3* (b)*, Δf1fo#107* (c)*, ΔfabH#4* (d) *and ΔfabH#6* (e). Red arrows (b-e) indicate the location of the gene intentionally deleted. The insert image (c) is an enlarged segments of *uncE* region, shown as a black box.

**Supplementary Fig. S5.**
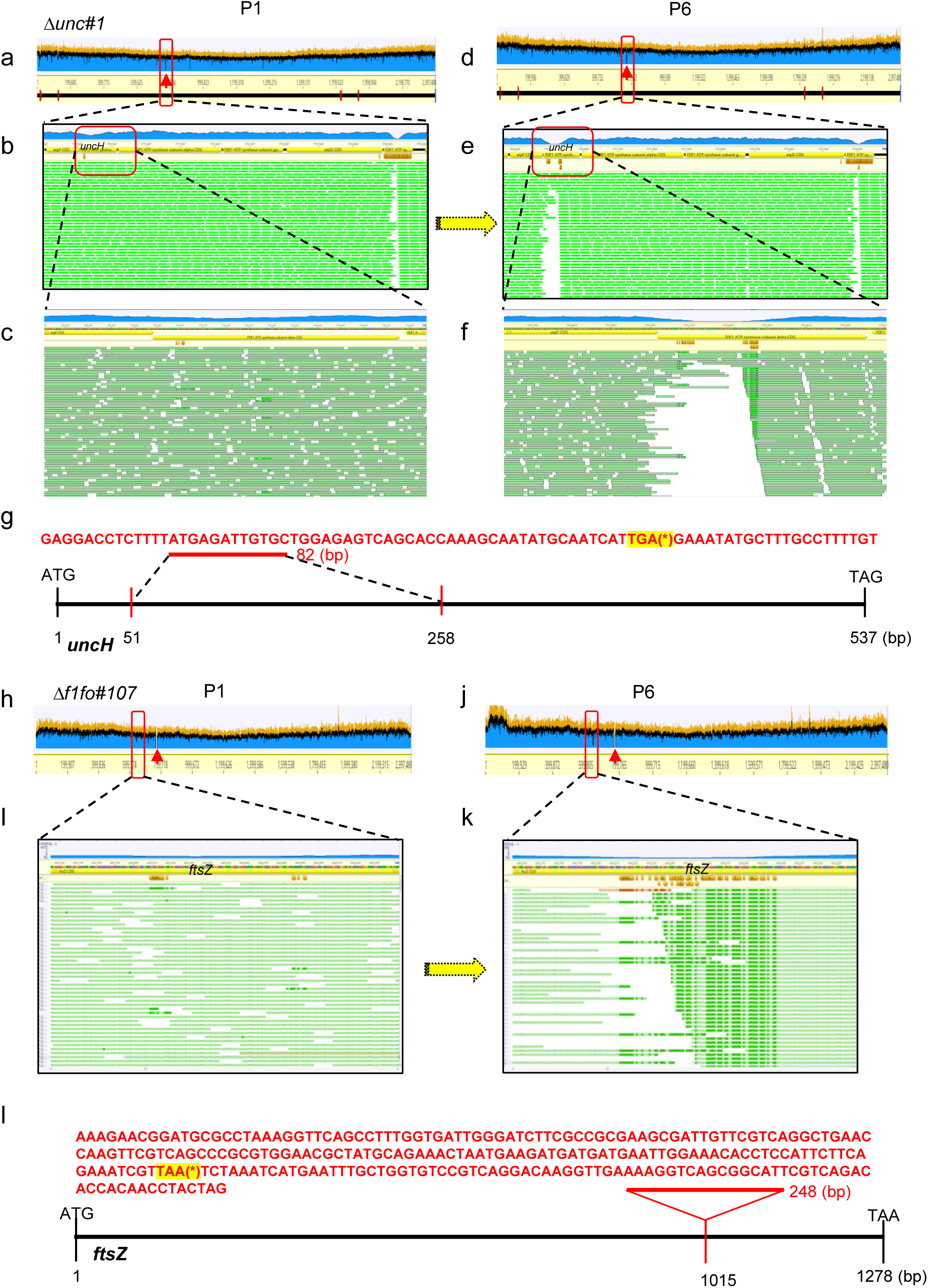
Appearance of mutations in essential genes in the evolved populations of strains intentionally deleted for different essential genes. (a-f) Aligned sequence reads from *ΔuncC#1* of P1 population (a-c) and P6 population (d-f). Red arrows indicate the deleted *uncC* in the genome (a and d) in the *ΔuncC#1* strain. Red box indicates mutation of *uncH* in *ΔuncC#1* (a and d). Enlarged mutated *uncH* region of aligned sequence reads in *ΔuncC#1* of P1 population (b-c) and P6 population (e-f). The yellow arrow indicates the passage procedure. (g) Sequencing of the mutated *uncH* gene in *ΔuncC#1* in P6. Red letters indicate a fragment (82 bp) that was present in mutated *uncH* gene (upper). Asterisk (*) indicates the stop codon that was highlighted. Vertical bars indicate the coordinates of the *uncH* ORF (lower). The coordinates between two red bars indicate replaced region. (h-k) Aligned sequence reads from *Δf1fo#107* of P1 population (h-i) and P6 population (j-k). Red arrows indicate the deleted *f1fo* in the genome (h and j) in the *Δf1fo#107*. Red box indicates mutation of *ftsZ* in *Δf1fo#107* (h and j). Enlarged mutated *ftsZ* region of aligned sequence reads in *Δf1fo#107* of P1 population (h) and P6 population (i). The yellow arrow indicates the passage procedure. Supplementary Fig.5J is a duplicate of Supplementary Fig.4B to show a genome alignment of *Δf1fo#107* of P6. (l) Sequencing of the mutated *ftsZ* in *Δf1fo#107* in P6. Red letters indicate a fragment (248 bp) that was present in mutated *Δf1fo#107* (upper). Asterisk (*) indicates the stop codon that was highlighted. Vertical bars indicated the coordinates of the *uncH* ORF (lower). The red bar indicates the coordinate where insertion occurred.

**Supplementary Fig. 6.**
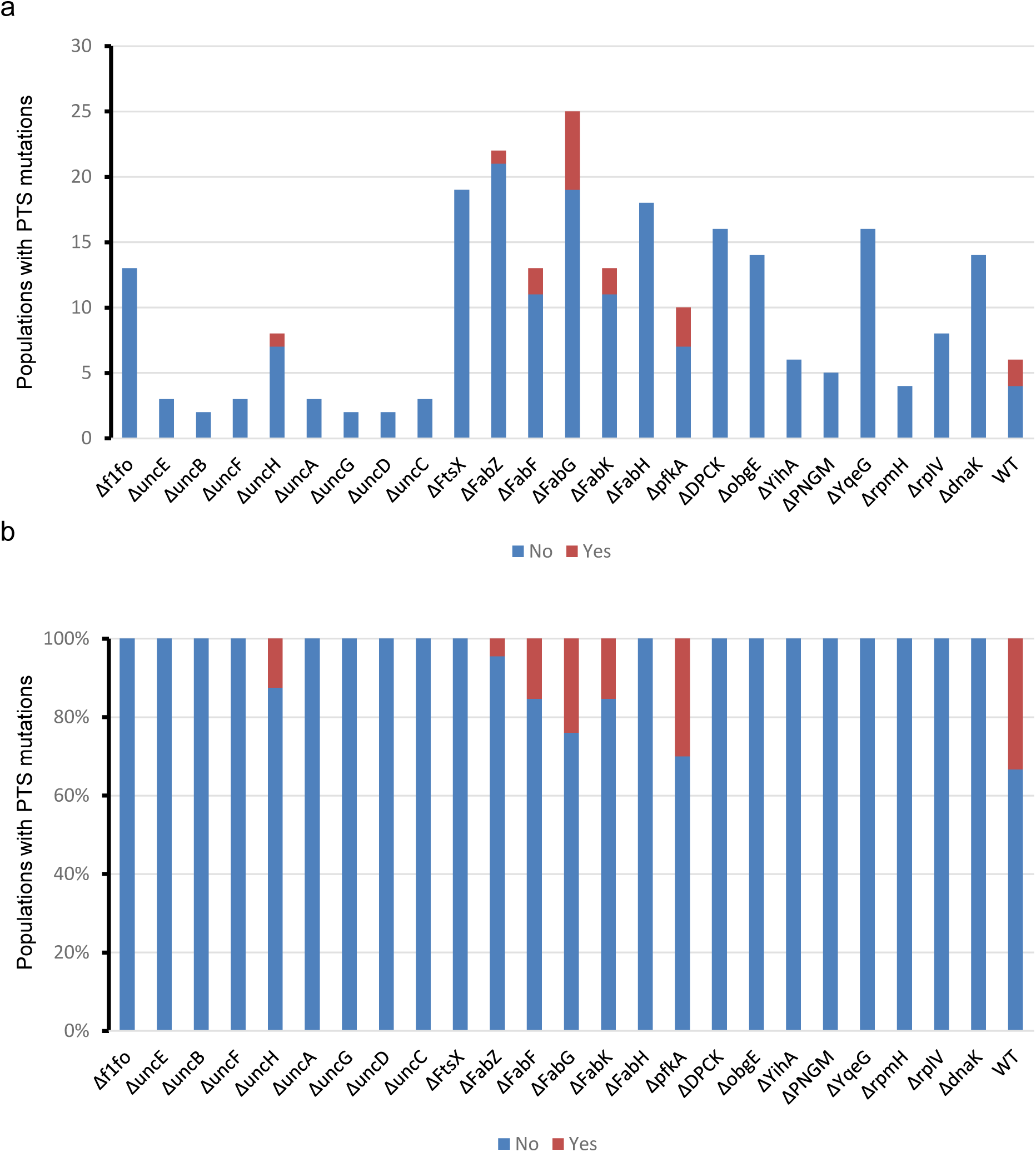
Populations with and without mutations in three PTS mannose/fructose/sorbose transporter subunits. (a-b) Number (a) and percentage (b) of populations that have mutations in any of three PTS mannose/fructose/sorbose transporter subunits, namely J1C87_09355, J1C87_09360, and J1C87_09365.

**Supplementary Fig. 7.**
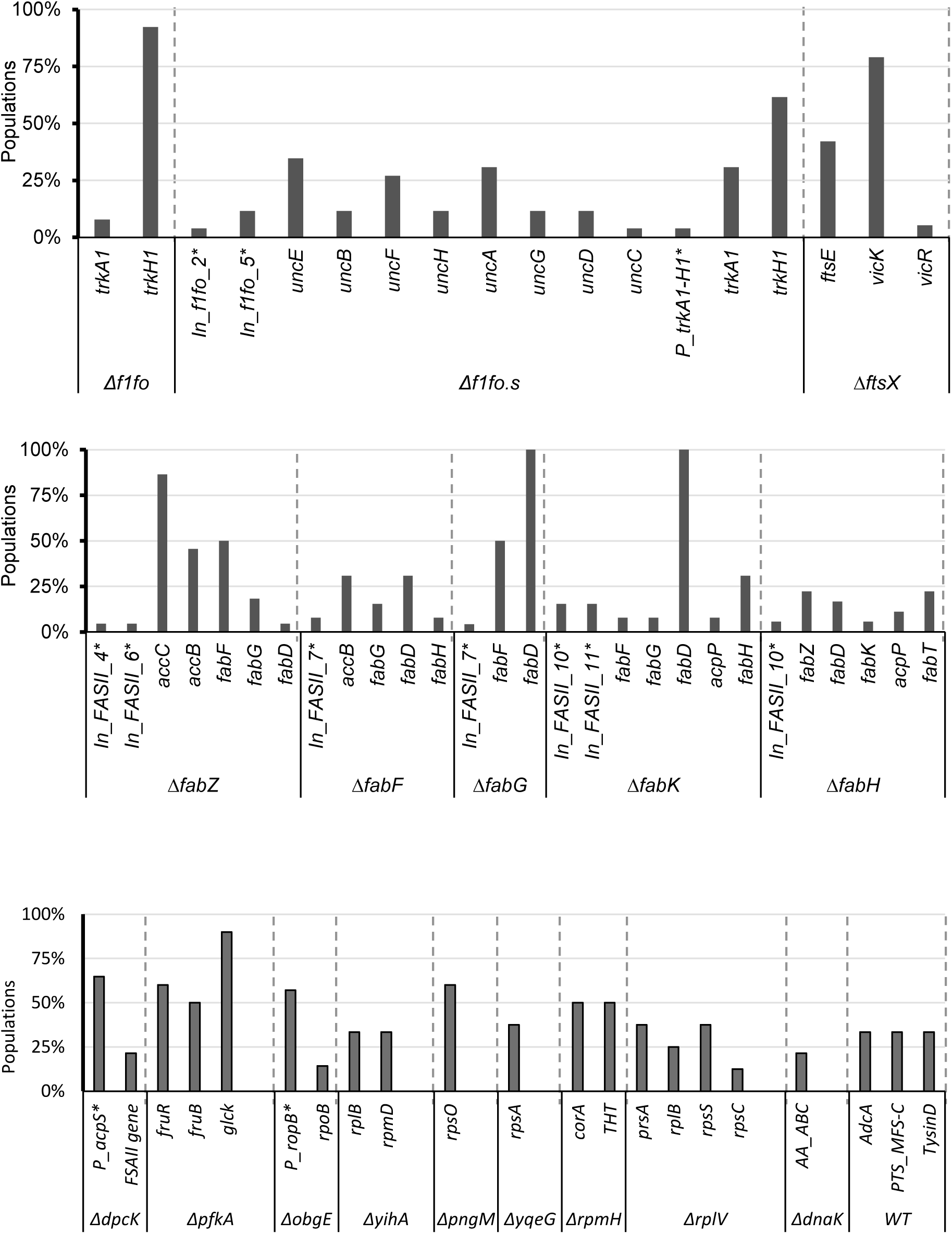
The most frequent suppressors identified in mutants deleted of essential genes. The percentage of mutated segments in evolved populations in *Δf1fo*, mutants deleted of single *unc* subunit genes, denoted as *Δf1fo.s*, the other single mutants indicated, and WT. The names with an asterisk (*) indicate the intergenic regions.

**Supplementary Fig. 8.**
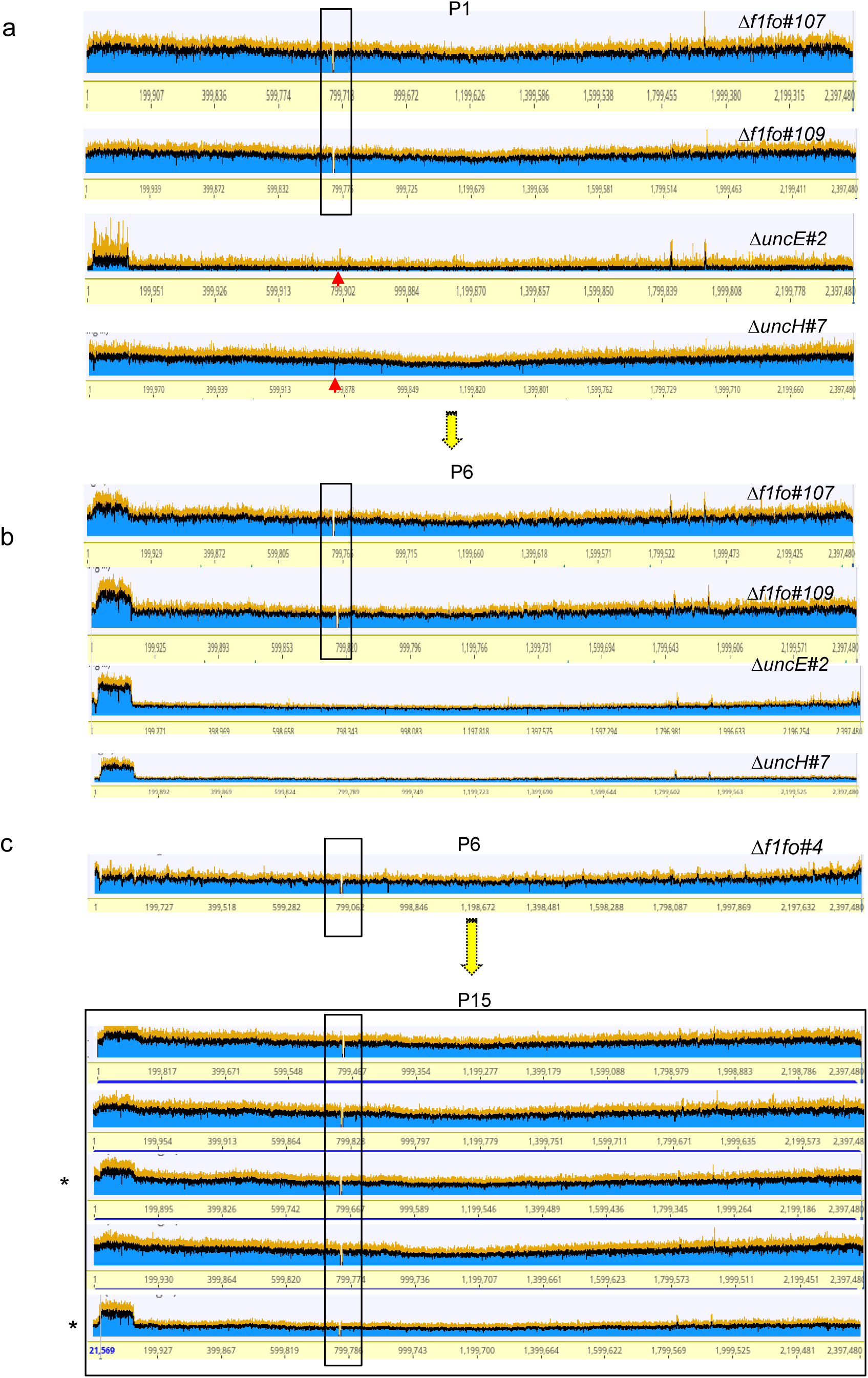
Appearance of *v1vo* region gene duplication during passage. (Related to Figure 4a and 4b) (a-b) Alignment of sequence reads in populations of *Δf1fo#107, Δf1fo#109, uncE#2* and *uncH#7* at P1 (A) and P6 (B). The black box indicates the deleted *f1fo* region and the red arrows indicate the deleted *uncE* or *uncH*. (c) Alignment of sequence reads of the P6 population in *Δf1fo#4* (upper) and five derived populations from P15 (lower). Rectangles represent the deleted *f1fo* region. Asterisks (*) indicate the two populations with *v1vo* region gene duplications.

**Supplementary Fig. 9.**
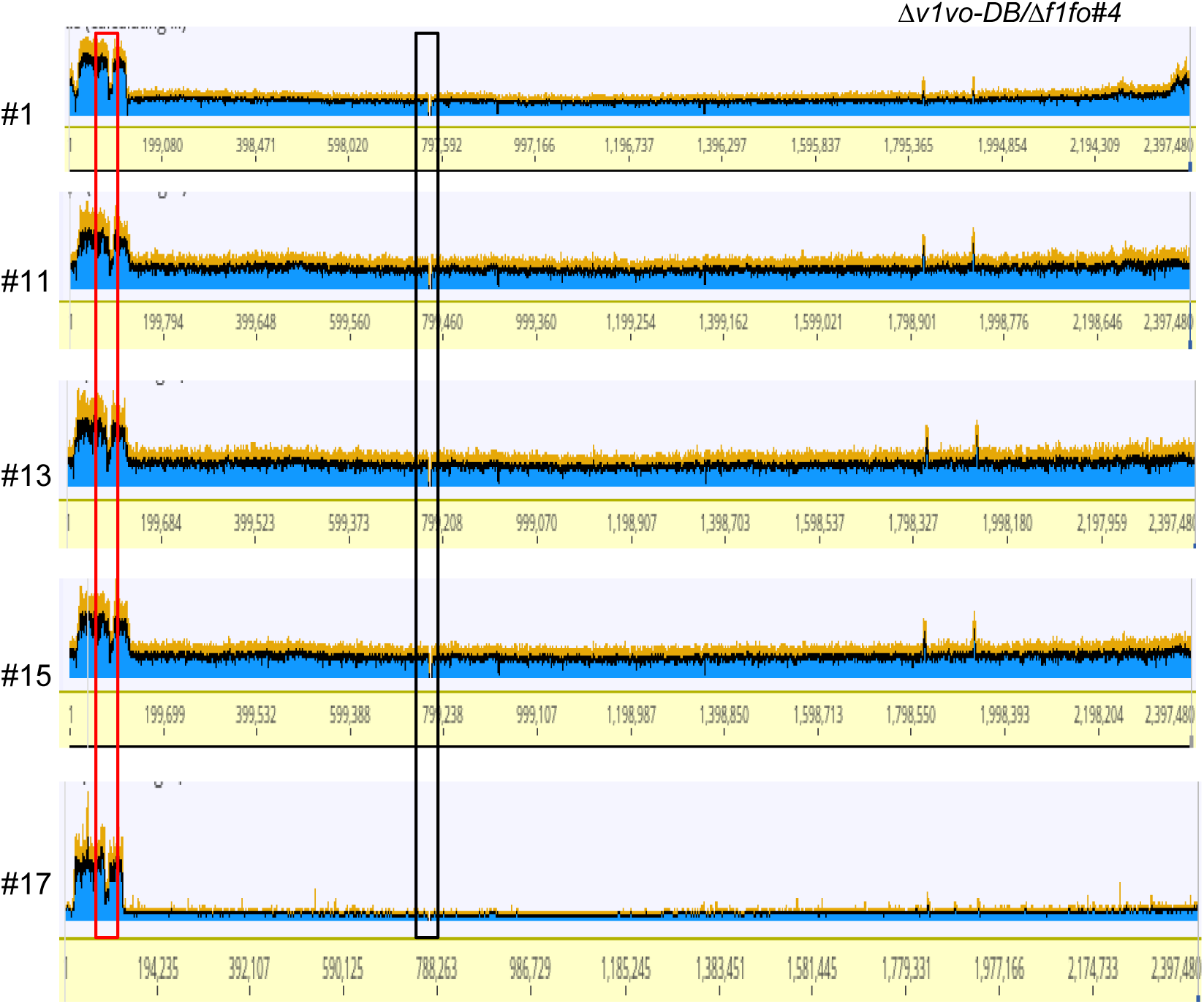
Gene duplication in *Δv1vo-DB/Δf1fo#4* populations (related to Figure 4c). Alignment of sequence reads of five additional *Δv1vo-DB/Δf1fo#4* populations. The black box indicates the deleted *f1fo*. The red box indicates the gene duplication of *v1vo* region when one copy of *v1vo* was deleted.

**Supplementary Fig. 10.**
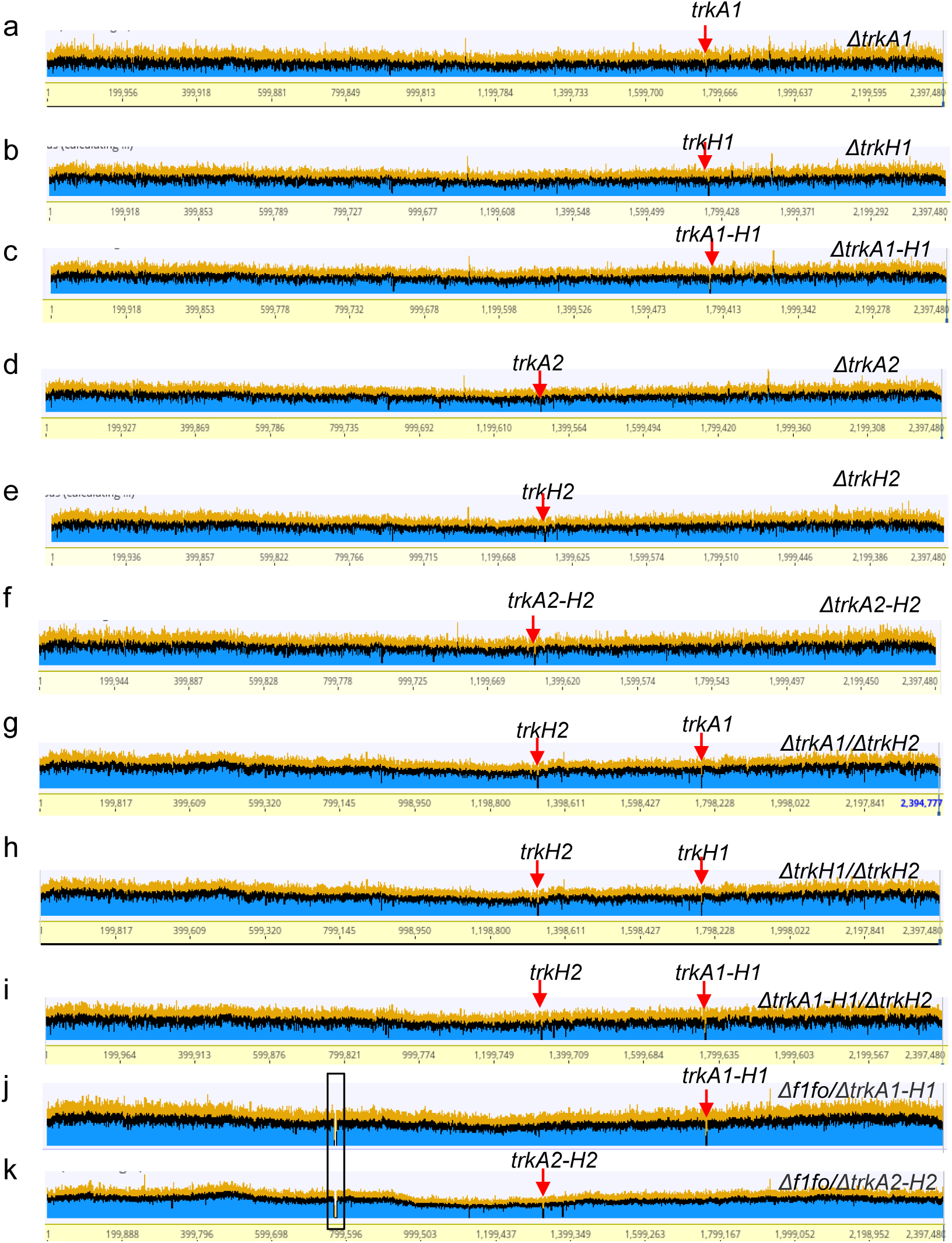
Alignment of sequence reads of *trk* mutants with and without *f1fo* deletion. (a-k) Alignment of sequence reads of *ΔtrkA1* (a), *ΔtrkH1* (b), *ΔtrkA1-H1* (c), *ΔtrkA2* (d), *ΔtrkH2* (e), *ΔtrkA2-H2* (f), *ΔtrkA1/ΔtrkH2* (g), *ΔtrkH1/ΔtrkH2* (h), *ΔtrkA1-H1/ΔtrkH2* (i), *Δf1fo/ΔtrkA1-H1* (j), and *Δf1fo/ΔtrkA2-H2* (k). Red arrows indicated intended gene deletion. The block box indicates the deleted *f1fo region.* (related to Figure 4f).

**Supplementary Fig. 11.**
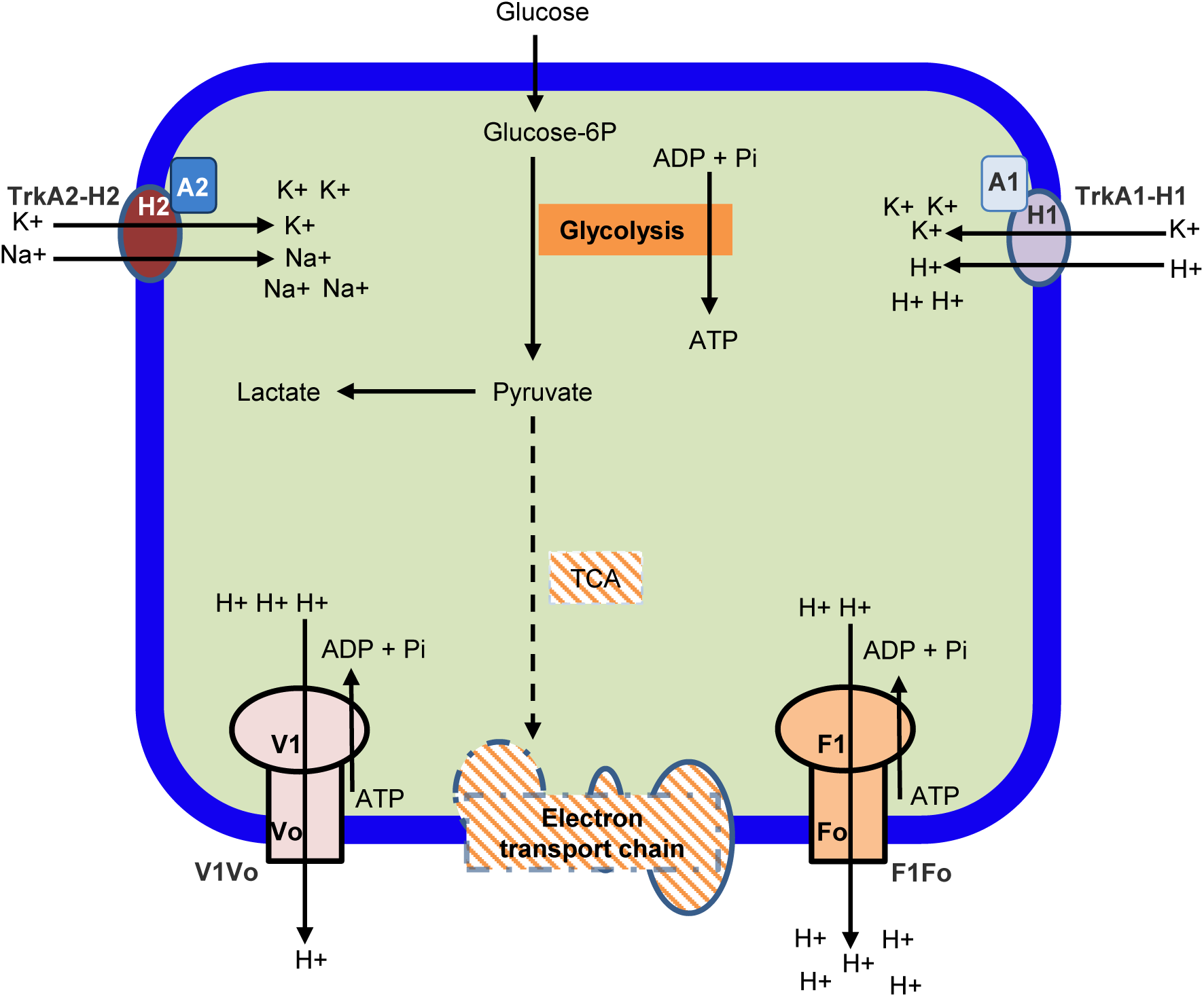
Working model for F1Fo, V1Vo, TrkA1-H1 and TrkA2-H2. For streptococci, which do not contain a complete tricarboxylic acid cycle (TCA: shaded) or electron transport chain (ETC: shaded), energy production is mainly through lactic acid fermentation. The lactate from fermentation creates acidic conditions in the cytosol, which becomes toxic if H+ is not exported. F1Fo pumps out H+ from the cytosol using ATP, and V1Vo supplements or replaces the function of F1Fo when the F1Fo is inhibited/inactivated. Both TrkA1-H1 and TrkA2-H1 are potassium transporters, and functionally redundant. TrkA1-H1 pumps in potassium together with H+, acidifying the cytosol when H+ is not exported. In contrast, TrkA2-H2 does not pump protons into the cytosol when pumping in potassium (likely instead pumping in Na+). Therefore, the function of TrkA1-H1 must be inhibited if F1Fo is not functional. Solid lines indicate presence of reactions. Dashed lines indicate absence of reactions.

**Supplementary Fig. 12.**
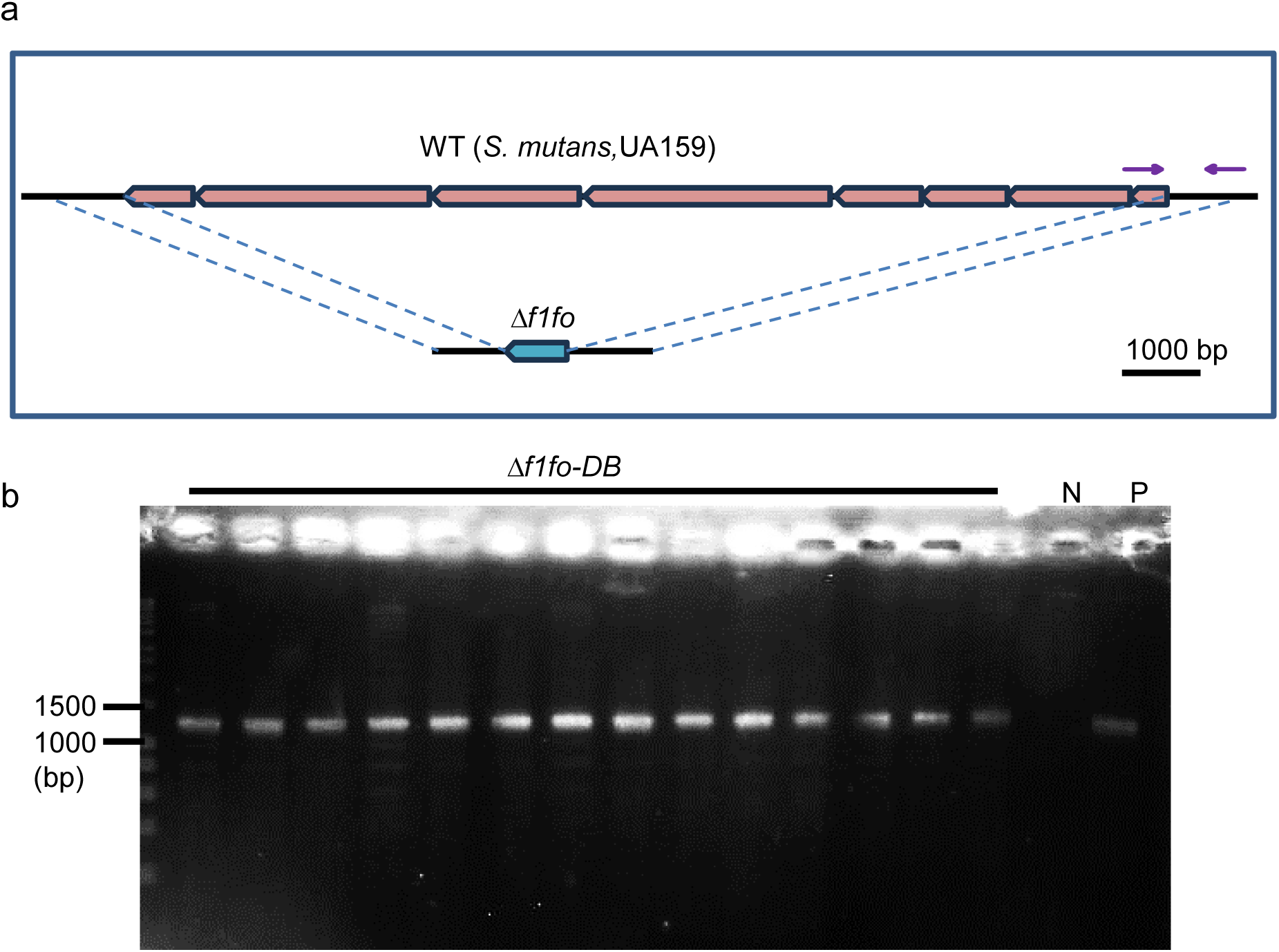
*f1fo* cannot be deleted in *S. mutans* UA159. (a) Gene knockout strategy to delete entire region of eight *f1fo* subunits by homologous recombination. The purple blocks indicate the eight F1Fo subunit genes in WT (upper). The blue block indicates the *kan* gene in *Δf1fo* mutants (lower). The two arrows indicate the primer set for the PCR amplicon of the region in (B). (b) Genotyping the presence of wild-type *f1fo* colonies from selection medium. P: positive control using DNA template from wild-type. N: negative control using water as a template. The size of amplicons is about 1300 bp.

**Supplementary Fig. 13.**
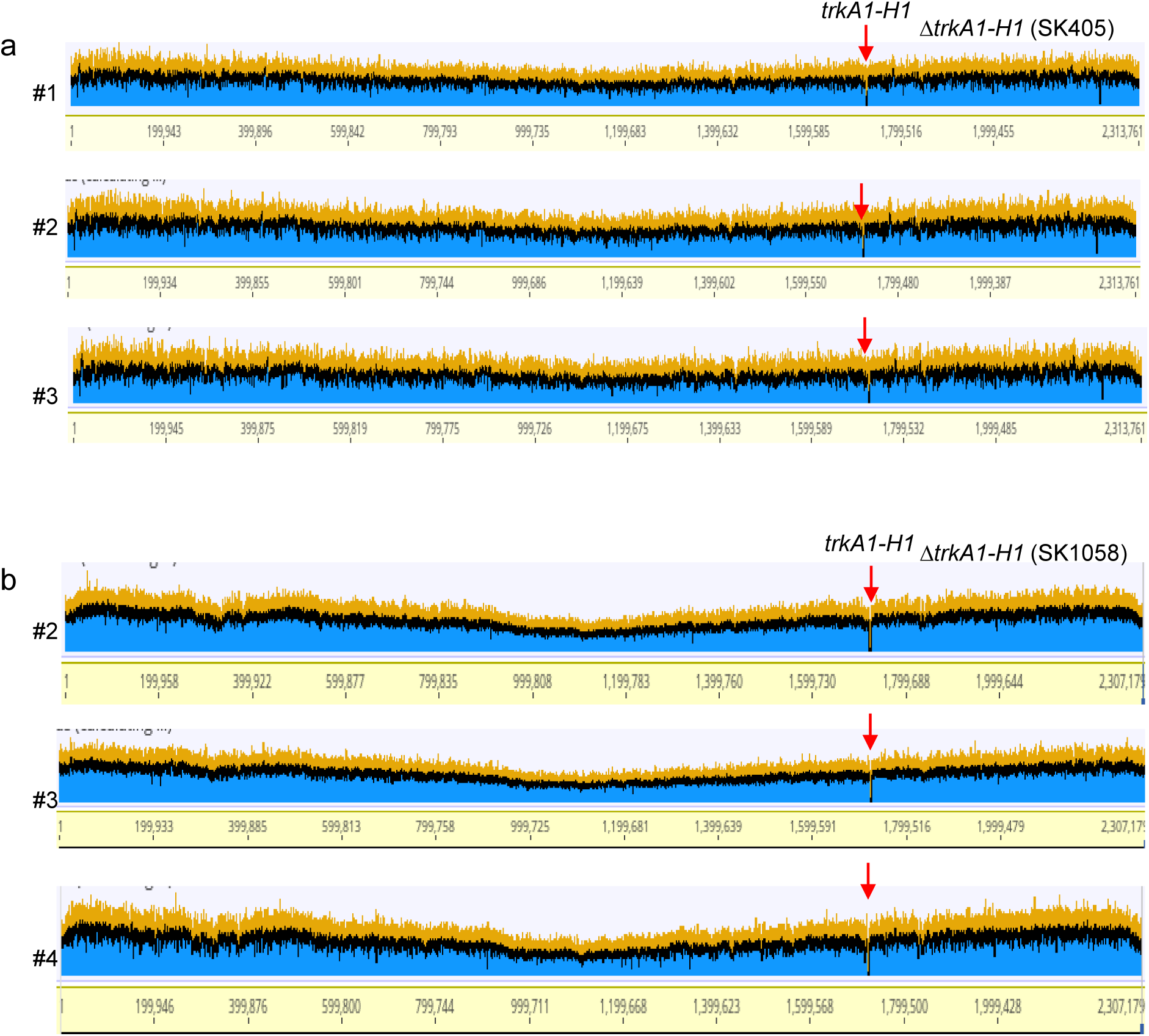
Deletion *of trkA1-H1* in *S. sanguinis* SK1058 and S. sanguinis SK405 (related to Figure 5c). (a-b) Alignment of sequence reads of *ΔtrkA1-H1* populations in SK405 (a) and in SK1058 (b) backgrounds.

**Supplementary Fig. 14.**
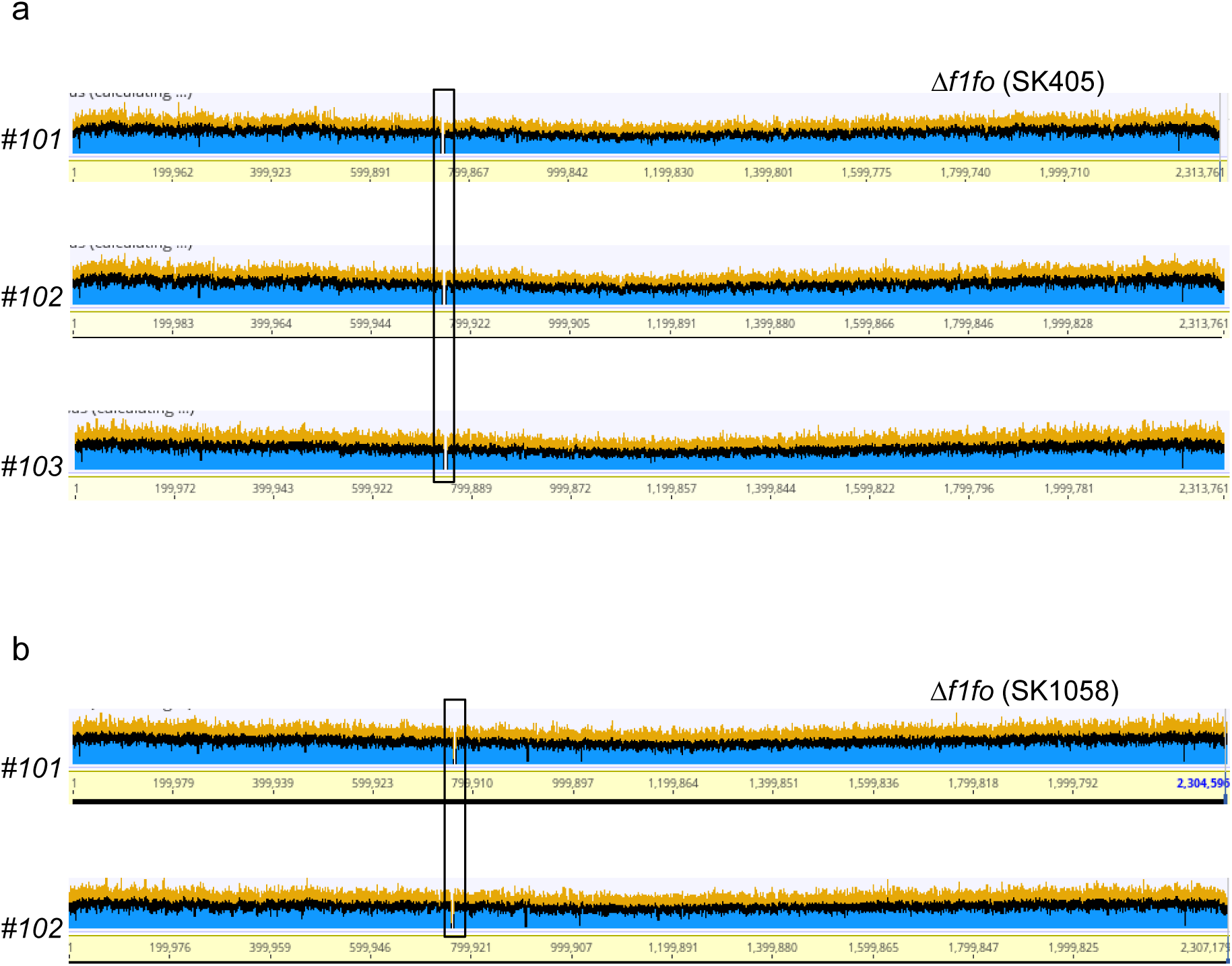
Deletion *of f1fo* in *S. sanguinis* SK1058 and SK405 (related to Figure 5c). Alignment of sequence reads of three *Δf1fo* populations in SK405 backgrounds (a) and two in SK1058 backgrounds (b) (related to Figure 5c). Rectangles represent the deleted *f1fo* region.

## References

1. Xu, P., Ge, X., Chen, L., Wang, X., Dou, Y., Xu, J.Z., Patel, J.R., Stone, V., Trinh, M., Evans, K., et al. (2011). Genome-wide essential gene identification in *Streptococcus sanguinis*. Scientific reports 1, 125. 10.1038/srep00125.

2. Rancati, G., Moffat, J., Typas, A., and Pavelka, N. (2018). Emerging and evolving concepts in gene essentiality. Nat Rev Genet 19, 34–49. 10.1038/nrg.2017.74.

3. Zhang, Z., and Ren, Q. (2015). Why are essential genes essential? - The essentiality of *Saccharomyces genes*. Microb Cell 2, 280–287. 10.15698/mic2015.08.218.

4. D’Elia, M.A., Pereira, M.P., and Brown, E.D. (2009). Are essential genes really essential? Trends Microbiol 17, 433–438. 10.1016/j.tim.2009.08.005.

5. Brown, E.D., and Wright, G.D. (2016). Antibacterial drug discovery in the resistance era. Nature 529, 336–343. 10.1038/nature17042.

6. Hogan, A.M., and Cardona, S.T. (2022). Gradients in gene essentiality reshape antibacterial research. FEMS Microbiol Rev 46. 10.1093/femsre/fuac005.

7. Costanzo, M., VanderSluis, B., Koch, E.N., Baryshnikova, A., Pons, C., Tan, G., Wang, W., Usaj, M., Hanchard, J., Lee, S.D., et al. (2016). A global genetic interaction network maps a wiring diagram of cellular function. Science 353. 10.1126/science.aaf1420.

8. Jana, B., Liu, X., Dénéréaz, J., Park, H., Leshchiner, D., Liu, B., Gallay, C., Veening, J.W., and van Opijnen, T. (2023). CRISPRi-TnSeq: A genome-wide high-throughput tool for bacterial essential-nonessential genetic interaction mapping. bioRxiv. 10.1101/2023.05.31.543074.

9. Nijman, S.M. (2011). Synthetic lethality: general principles, utility and detection using genetic screens in human cells. FEBS Lett 585, 1–6. 10.1016/j.febslet.2010.11.024.

10. Leaver, M., Domínguez-Cuevas, P., Coxhead, J.M., Daniel, R.A., and Errington, J. (2009). Life without a wall or division machine in *Bacillus subtilis*. Nature 457, 849–853. 10.1038/nature07742.

11. Mutschler, H., Gebhardt, M., Shoeman, R.L., and Meinhart, A. (2011). A novel mechanism of programmed cell death in bacteria by toxin-antitoxin systems corrupts peptidoglycan synthesis. PLoS Biol 9, e1001033. 10.1371/journal.pbio.1001033.

12. Li, J., Wang, H.T., Wang, W.T., Zhang, X.R., Suo, F., Ren, J.Y., Bi, Y., Xue, Y.X., Hu, W., Dong, M.Q., and Du, L.L. (2019). Systematic analysis reveals the prevalence and principles of bypassable gene essentiality. Nat Commun 10, 1002. 10.1038/s41467-019-08928-1.

13. Larrimore, K.E., and Rancati, G. (2019). The conditional nature of gene essentiality in *Pseudomonas aeruginosa*. Curr Opin Genet Dev 58-59, 55-61. 10.1016/j.gde.2019.07.015.

14. Liu, G., Yong, M.Y., Yurieva, M., Srinivasan, K.G., Liu, J., Lim, J.S., Poidinger, M., Wright, G.D., Zolezzi, F., Choi, H., et al. (2015). Gene Essentiality Is a Quantitative Property Linked to Cellular Evolvability. Cell 163, 1388–1399. 10.1016/j.cell.2015.10.069.

15. Rancati, G., Pavelka, N., Fleharty, B., Noll, A., Trimble, R., Walton, K., Perera, A., Staehling-Hampton, K., Seidel, C.W., and Li, R. (2008). Aneuploidy underlies rapid adaptive evolution of yeast cells deprived of a conserved cytokinesis motor. Cell 135, 879–893. 10.1016/j.cell.2008.09.039.

16. Bao, L., Inoue, N., Ishikawa, M., Gotoh, E., Teh, O.K., Higa, T., Morimoto, T., Ginanjar, E.F., Harashima, H., Noda, N., et al. (2022). A PSTAIRE-type cyclin-dependent kinase controls light responses in land plants. Sci Adv 8, eabk2116. 10.1126/sciadv.abk2116.

17. Rosconi, F., Rudmann, E., Li, J., Surujon, D., Anthony, J., Frank, M., Jones, D.S., Rock, C., Rosch, J.W., Johnston, C.D., and van Opijnen, T. (2022). A bacterial pan-genome makes gene essentiality strain-dependent and evolvable. Nat Microbiol 7, 1580–1592. 10.1038/s41564-022-01208-7.

18. Couce, A., Limdi, A., Magnan, M., Owen, S.V., Herren, C.M., Lenski, R.E., Tenaillon, O., and Baym, M. (2024). Changing fitness effects of mutations through long-term bacterial evolution. Science 383, eadd1417. 10.1126/science.add1417.

19. Santamaría, D., Barrière, C., Cerqueira, A., Hunt, S., Tardy, C., Newton, K., Cáceres, J.F., Dubus, P., Malumbres, M., and Barbacid, M. (2007). Cdk1 is sufficient to drive the mammalian cell cycle. Nature 448, 811–815. 10.1038/nature06046.

20. Nowack, M.K., Harashima, H., Dissmeyer, N., Zhao, X.A., Bouyer, D., Weimer, A.K., De Winter, F., Yang, F., and Schnittger, A. (2012). Genetic framework of cyclin-dependent kinase function in *Arabidopsis*. Dev. Cell 22, 1030–1040. 10.1016/j.devcel.2012.02.015.

21. Glass, J.I., Merryman, C., Wise, K.S., Hutchison, C.A., and Smith, H.O. (2017). Minimal Cells-Real and Imagined. Cold Spring Harb Perspect Biol 9. 10.1101/cshperspect.a023861.

22. Juhas, M., Eberl, L., and Church, G.M. (2012). Essential genes as antimicrobial targets and cornerstones of synthetic biology. Trends Biotechnol 30, 601–607. 10.1016/j.tibtech.2012.08.002.

23. Hutchison, C.A., Chuang, R.Y., Noskov, V.N., Assad-Garcia, N., Deerinck, T.J., Ellisman, M.H., Gill, J., Kannan, K., Karas, B.J., Ma, L., et al. (2016). Design and synthesis of a minimal bacterial genome. Science 351, aad6253. 10.1126/science.aad6253.

24. Ratcliff, W.C., Denison, R.F., Borrello, M., and Travisano, M. (2012). Experimental evolution of multicellularity. Proc Natl Acad Sci U S A 109, 1595–1600. 10.1073/pnas.1115323109.

25. Blount, Z.D., Borland, C.Z., and Lenski, R.E. (2008). Historical contingency and the evolution of a key innovation in an experimental population of Escherichia coli. Proc Natl Acad Sci U S A 105, 7899–7906. 10.1073/pnas.0803151105.

26. Moger-Reischer, R.Z., Glass, J.I., Wise, K.S., Sun, L., Bittencourt, D.M.C., Lehmkuhl, B.K., Schoolmaster, D.R., Lynch, M., and Lennon, J.T. (2023). Evolution of a minimal cell. Nature 620, 122–127. 10.1038/s41586-023-06288-x.

27. Kawecki, T.J., Lenski, R.E., Ebert, D., Hollis, B., Olivieri, I., and Whitlock, M.C. (2012). Experimental evolution. Trends Ecol Evol 27, 547–560. 10.1016/j.tree.2012.06.001.

28. Barrick, J.E., and Lenski, R.E. (2013). Genome dynamics during experimental evolution. Nat Rev Genet 14, 827–839. 10.1038/nrg3564.

29. McDonald, M.J. (2019). Microbial Experimental Evolution - a proving ground for evolutionary theory and a tool for discovery. EMBO Rep 20, e46992. 10.15252/embr.201846992.

30. Van den Bergh, B., Swings, T., Fauvart, M., and Michiels, J. (2018). Experimental Design, Population Dynamics, and Diversity in Microbial Experimental Evolution. Microbiol Mol Biol Rev 82. 10.1128/MMBR.00008-18.

31. Taylor, T.B., Mulley, G., Dills, A.H., Alsohim, A.S., McGuffin, L.J., Studholme, D.J., Silby, M.W., Brockhurst, M.A., Johnson, L.J., and Jackson, R.W. (2015). Evolutionary resurrection of flagellar motility via rewiring of the nitrogen regulation system. Science 347, 1014–1017. 10.1126/science.1259145.

32. Xu, P., Alves, J.M., Kitten, T., Brown, A., Chen, Z., Ozaki, L.S., Manque, P., Ge, X., Serrano, M.G., Puiu, D., et al. (2007). Genome of the opportunistic pathogen Streptococcus sanguinis. J Bacteriol 189, 3166–3175. 10.1128/JB.01808-06.

33. Lázár, V., Pal Singh, G., Spohn, R., Nagy, I., Horváth, B., Hrtyan, M., Busa-Fekete, R., Bogos, B., Méhi, O., Csörgő, B., et al. (2013). Bacterial evolution of antibiotic hypersensitivity. Mol Syst Biol 9, 700. 10.1038/msb.2013.57.

34. Rhodes, D.V., Crump, K.E., Makhlynets, O., Snyder, M., Ge, X., Xu, P., Stubbe, J., and Kitten, T. (2014). Genetic characterization and role in virulence of the ribonucleotide reductases of Streptococcus sanguinis. J Biol Chem 289, 6273–6287. 10.1074/jbc.M113.533620.

35. Luo, H., Lin, Y., Gao, F., Zhang, C.T., and Zhang, R. (2014). DEG 10, an update of the database of essential genes that includes both protein-coding genes and noncoding genomic elements. Nucleic Acids Res 42, D574–580. 10.1093/nar/gkt1131.

36. Nelson, N., Perzov, N., Cohen, A., Hagai, K., Padler, V., and Nelson, H. (2000). The cellular biology of proton-motive force generation by V-ATPases. J Exp Biol 203, 89–95. 10.1242/jeb.203.1.89.

37. Röttig, A., and Steinbüchel, A. (2013). Acyltransferases in bacteria. Microbiol Mol Biol Rev 77, 277–321. 10.1128/MMBR.00010-13.

38. Baker, S.P., Nulton, T.J., and Kitten, T. (2019). Genomic, Phenotypic, and Virulence Analysis of Streptococcus sanguinis Oral and Infective-Endocarditis Isolates. Infect Immun 87. 10.1128/IAI.00703-18.

39. Ghosh, A., N, S., and Saha, S. (2020). Survey of drug resistance associated gene mutations in Mycobacterium tuberculosis, ESKAPE and other bacterial species. Scientific reports 10, 8957. 10.1038/s41598-020-65766-8.

40. Zeng, L., Walker, A.R., Lee, K., Taylor, Z.A., and Burne, R.A. (2021). Spontaneous Mutants of Streptococcus sanguinis with Defects in the Glucose-Phosphotransferase System Show Enhanced Post-Exponential-Phase Fitness. J Bacteriol 203, e0037521. 10.1128/JB.00375-21.

41. Chevin, L.M., Martin, G., and Lenormand, T. (2010). Fisher’s model and the genomics of adaptation: restricted pleiotropy, heterogenous mutation, and parallel evolution. Evolution 64, 3213–3231. 10.1111/j.1558-5646.2010.01058.x.

42. Wichman, H.A., Badgett, M.R., Scott, L.A., Boulianne, C.M., and Bull, J.J. (1999). Different trajectories of parallel evolution during viral adaptation. Science 285, 422–424. 10.1126/science.285.5426.422.

43. Stern, D.L. (2013). The genetic causes of convergent evolution. Nat Rev Genet 14, 751–764. 10.1038/nrg3483.

44. Sheng, J., and Marquis, R.E. (2006). Enhanced acid resistance of oral streptococci at lethal pH values associated with acid-tolerant catabolism and with ATP synthase activity. FEMS Microbiol Lett 262, 93–98. 10.1111/j.1574-6968.2006.00374.x.

45. Corratgé-Faillie, C., Jabnoune, M., Zimmermann, S., Véry, A.A., Fizames, C., and Sentenac, H. (2010). Potassium and sodium transport in non-animal cells: the Trk/Ktr/HKT transporter family. Cell Mol Life Sci 67, 2511–2532. 10.1007/s00018-010-0317-7.

46. Du, S., Pichoff, S., and Lutkenhaus, J. (2016). FtsEX acts on FtsA to regulate divisome assembly and activity. Proc Natl Acad Sci U S A 113, E5052–5061. 10.1073/pnas.1606656113.

47. Moraes, J.J., Stipp, R.N., Harth-Chu, E.N., Camargo, T.M., Höfling, J.F., and Mattos-Graner, R.O. (2014). Two-component system VicRK regulates functions associated with establishment of Streptococcus sanguinis in biofilms. Infect Immun 82, 4941–4951. 10.1128/IAI.01850-14.

48. Sham, L.T., Barendt, S.M., Kopecky, K.E., and Winkler, M.E. (2011). Essential PcsB putative peptidoglycan hydrolase interacts with the essential FtsXSpn cell division protein in *Streptococcus pneumoniae* D39. Proc Natl Acad Sci U S A 108, E1061–1069. 10.1073/pnas.1108323108.

49. Gupta, A., Singh, P.K., Sharma, P., Kaur, P., Sharma, S., and Singh, T.P. (2020). Structural and biochemical studies of phosphopantetheine adenylyltransferase from *Acinetobacter baumannii* with dephospho-coenzyme A and coenzyme A. Int J Biol Macromol 142, 181–190. 10.1016/j.ijbiomac.2019.09.090.

50. Heath, R.J., and Rock, C.O. (1996). Inhibition of beta-ketoacyl-acyl carrier protein synthase III (FabH) by acyl-acyl carrier protein in Escherichia coli. The Journal of biological chemistry 271, 10996–11000. 10.1074/jbc.271.18.10996.

51. Phong, W.Y., Lin, W., Rao, S.P., Dick, T., Alonso, S., and Pethe, K. (2013). Characterization of phosphofructokinase activity in *Mycobacterium tuberculosis* reveals that a functional glycolytic carbon flow is necessary to limit the accumulation of toxic metabolic intermediates under hypoxia. PLoS One 8, e56037. 10.1371/journal.pone.0056037.

52. Skarlatos, P., and Dahl, M.K. (1998). The glucose kinase of Bacillus subtilis. J Bacteriol 180, 3222–3226. 10.1128/JB.180.12.3222-3226.1998.

53. Boulanger, E.F., Sabag-Daigle, A., Thirugnanasambantham, P., Gopalan, V., and Ahmer, B.M.M. (2021). Sugar-Phosphate Toxicities. Microbiol Mol Biol Rev 85, e0012321. 10.1128/MMBR.00123-21.

54. Zeng, L., and Burne, R.A. (2021). Molecular mechanisms controlling fructose-specific memory and catabolite repression in lactose metabolism by Streptococcus mutans. Mol Microbiol 115, 70–83. 10.1111/mmi.14597.

55. Morrissey, A.T., and Fraenkel, D.G. (1972). Suppressor of phosphofructokinase mutations of Escherichia coli. J Bacteriol 112, 183–187. 10.1128/jb.112.1.183-187.1972.

56. Babul, J. (1978). Phosphofructokinases from Escherichia coli. Purification and characterization of the nonallosteric isozyme. The Journal of biological chemistry 253, 4350–4355.

57. Deckers, B., Vercauteren, S., De Kock, V., Martin, C., Lazar, T., Herpels, P., Dewachter, L., Verstraeten, N., Peeters, E., Ballet, S., et al. (2023). YbiB: a novel interactor of the GTPase ObgE. Nucleic Acids Res 51, 3420–3435. 10.1093/nar/gkad127.

58. Vidwans, S.J., Ireton, K., and Grossman, A.D. (1995). Possible role for the essential GTP-binding protein Obg in regulating the initiation of sporulation in Bacillus subtilis. J Bacteriol 177, 3308–3311. 10.1128/jb.177.11.3308-3311.1995.

59. Okamoto, S., Itoh, M., and Ochi, K. (1997). Molecular cloning and characterization of the obg gene of Streptomyces griseus in relation to the onset of morphological differentiation. J Bacteriol 179, 170–179. 10.1128/jb.179.1.170-179.1997.

60. Okamoto, S., and Ochi, K. (1998). An essential GTP-binding protein functions as a regulator for differentiation in Streptomyces coelicolor. Mol Microbiol 30, 107–119. 10.1046/j.1365-2958.1998.01042.x.

61. Verstraeten, N., Knapen, W.J., Kint, C.I., Liebens, V., Van den Bergh, B., Dewachter, L., Michiels, J.E., Fu, Q., David, C.C., Fierro, A.C., et al. (2015). Obg and Membrane Depolarization Are Part of a Microbial Bet-Hedging Strategy that Leads to Antibiotic Tolerance. Molecular cell 59, 9–21. 10.1016/j.molcel.2015.05.011.

62. Feng, B., Mandava, C.S., Guo, Q., Wang, J., Cao, W., Li, N., Zhang, Y., Wang, Z., Wu, J., Sanyal, S., et al. (2014). Structural and functional insights into the mode of action of a universally conserved Obg GTPase. PLoS Biol 12, e1001866. 10.1371/journal.pbio.1001866.

63. Murti, K.G., Webster, R.G., and Jones, I.M. (1988). Localization of RNA polymerases on influenza viral ribonucleoproteins by immunogold labeling. Virology 164, 562–566. 10.1016/0042-6822(88)90574-0.

64. Zielke, R., Sikora, A., Dutkiewicz, R., Wegrzyn, G., and Czyż, A. (2003). Involvement of the cgtA gene function in stimulation of DNA repair in Escherichia coli and Vibrio harveyi. Microbiology (Reading) 149, 1763–1770. 10.1099/mic.0.26292-0.

65. Czyz, A., Zielke, R., Konopa, G., and Wegrzyn, G. (2001). A Vibrio harveyi insertional mutant in the cgtA (obg, yhbZ) gene, whose homologues are present in diverse organisms ranging from bacteria to humans and are essential genes in many bacterial species. Microbiology (Reading) 147, 183–191. 10.1099/00221287-147-1-183.

66. Courcelle, J., Khodursky, A., Peter, B., Brown, P.O., and Hanawalt, P.C. (2001). Comparative gene expression profiles following UV exposure in wild-type and SOS-deficient Escherichia coli. Genetics 158, 41–64. 10.1093/genetics/158.1.41.

67. Foti, J.J., Persky, N.S., Ferullo, D.J., and Lovett, S.T. (2007). Chromosome segregation control by Escherichia coli ObgE GTPase. Mol Microbiol 65, 569–581. 10.1111/j.1365-2958.2007.05811.x.

68. Tan, J., Jakob, U., and Bardwell, J.C. (2002). Overexpression of two different GTPases rescues a null mutation in a heat-induced rRNA methyltransferase. J Bacteriol 184, 2692–2698. 10.1128/JB.184.10.2692-2698.2002.

69. Nirody, J.A., Budin, I., and Rangamani, P. (2020). ATP synthase: Evolution, energetics, and membrane interactions. J Gen Physiol 152. 10.1085/jgp.201912475.

70. Lund, P., Tramonti, A., and De Biase, D. (2014). Coping with low pH: molecular strategies in neutralophilic bacteria. FEMS Microbiol Rev 38, 1091–1125. 10.1111/1574-6976.12076.

71. Wang, Y., Wu, J., Lv, M., Shao, Z., Hungwe, M., Wang, J., Bai, X., Xie, J., and Geng, W. (2021). Metabolism Characteristics of Lactic Acid Bacteria and the Expanding Applications in Food Industry. Front Bioeng Biotechnol 9, 612285. 10.3389/fbioe.2021.612285.

72. Ajdic, D., McShan, W.M., McLaughlin, R.E., Savic, G., Chang, J., Carson, M.B., Primeaux, C., Tian, R., Kenton, S., Jia, H., et al. (2002). Genome sequence of Streptococcus mutans UA159, a cariogenic dental pathogen. Proc Natl Acad Sci U S A 99, 14434–14439. 10.1073/pnas.172501299.

73. Chen, P., Wang, D., Chen, H., Zhou, Z., and He, X. (2016). The nonessentiality of essential genes in yeast provides therapeutic insights into a human disease. Genome Res 26, 1355–1362. 10.1101/gr.205955.116.

74. Mnaimneh, S., Davierwala, A.P., Haynes, J., Moffat, J., Peng, W.T., Zhang, W., Yang, X., Pootoolal, J., Chua, G., Lopez, A., et al. (2004). Exploration of essential gene functions via titratable promoter alleles. Cell 118, 31–44. 10.1016/j.cell.2004.06.013.

75. Peters, J.M., Colavin, A., Shi, H., Czarny, T.L., Larson, M.H., Wong, S., Hawkins, J.S., Lu, C.H.S., Koo, B.M., Marta, E., et al. (2016). A Comprehensive, CRISPR-based Functional Analysis of Essential Genes in Bacteria. Cell 165, 1493–1506. 10.1016/j.cell.2016.05.003.

76. Shields, R.C., Walker, A.R., Maricic, N., Chakraborty, B., Underhill, S.A.M., and Burne, R.A. (2020). Repurposing the Streptococcus mutans CRISPR-Cas9 System to Understand Essential Gene Function. PLoS Pathog 16, e1008344. 10.1371/journal.ppat.1008344.

77. Yother, J., Trieu-Cuot, P., Klaenhammer, T.R., and De Vos, W.M. (2002). Genetics of streptococci, lactococci, and enterococci: review of the sixth international conference. J Bacteriol 184, 6085–6092. 10.1128/JB.184.22.6085-6092.2002.

78. Otasek, D., Morris, J.H., Bouças, J., Pico, A.R., and Demchak, B. (2019). Cytoscape Automation: empowering workflow-based network analysis. Genome Biol 20, 185. 10.1186/s13059-019-1758-4.

